# *Porphyromonas gingivalis* activates Heat-Shock-Protein 27 to drive a LC3C-specific probacterial form of select autophagy that is redox sensitive for intracellular bacterial survival in human gingival mucosa

**DOI:** 10.1101/2024.07.01.601539

**Authors:** Bridgette Wellslager, JoAnn Roberts, Nityananda Chowdhury, Lalima Madan, Elsy Orellana, Özlem Yilmaz

## Abstract

*Porphyromonas gingivalis*, a major oral pathobiont, evades canonical host pathogen clearance in human primary gingival epithelial cells (GECs) by initiating a non-canonical variant of autophagy consisting of Microtubule-associated protein 1A/1B-light chain 3 (LC3)-rich autophagosomes, which then act as replicative niches. Simultaneously, *P. gingivalis* inhibits apoptosis and oxidative-stress, including extracellular-ATP (eATP)-mediated reactive-oxygen-species (ROS) production via phosphorylating Heat Shock Protein 27 (HSp27) with the bacterial nucleoside-diphosphate-kinase (Ndk). Here, we have mechanistically identified that *P. gingivalis*-mediated induction of HSp27 is crucial for the recruitment of the LC3 isoform, LC3C, to drive the formation of live *P. gingivalis*-containing Beclin1-ATG14-rich autophagosomes that are redox sensitive and non-degrading. HSp27 depletions of both infected GECs and gingiva-mimicking organotypic-culture systems resulted in the collapse of *P. gingivalis*-mediated autophagosomes, and abolished *P. gingivalis*-induced LC3C-specific autophagic-flux in a HSp27-dependent manner. Concurrently, HSp27 depletion accompanied by eATP treatment abrogated protracted Beclin 1-ATG14 partnering and decreased live intracellular *P. gingivalis* levels. These events were only partially restored via treatments with the antioxidant N-acetyl cysteine (NAC), which rescued the cellular redox environment independent of HSp27. Moreover, the temporal phosphorylation of HSp27 by the bacterial Ndk results in HSp27 tightly partnering with LC3C, hindering LC3C canonical cleavage, extending Beclin 1-ATG14 association, and halting canonical maturation. These findings pinpoint how HSp27 pleiotropically serves as a major platform-molecule, redox regulator, and stepwise modulator of LC3C during *P. gingivalis*-mediated non-canonical autophagy. Thus, our findings can determine specific molecular strategies for interfering with the host-adapted *P. gingivalis*’ successful mucosal colonization and oral dysbiosis.

## INTRODUCTION

*Porphyromonas gingivalis* is a Gram (-) fastidious obligate anaerobe that has been strongly linked to several debilitating chronic systemic diseases, including oro-digestive cancers, rheumatoid arthritis, and Alzheimer Disease [1–5]. *P. gingivalis* has been primarily studied as a prevailing pathogen in periodontal disease pathophysiology through the dysbiosis of the oral microbiota [5–9]. Recent studies, however, have underlined that live *P. gingivalis* is capable of invading the blood microvasculature of chronically inflamed gingival mucosa via an epithelial route, elucidating a systemic dissemination course and suggesting that the chronic colonization of *P. gingivalis* in the oral epithelia might act as a “seed bed” for the progression of *P. gingivalis-*associated chronic systemic diseases [10]. Human gingival epithelial cells (GECs) have been shown to act as replicative intracellular niches for *P. gingivalis,* despite otherwise acting as the first line of defense of the oral mucosa against numerous oral pathogens [7,11–15]. *P. gingivalis* is capable of invading traditionally defensive GECs, in a manner that is initially mediated by *P. gingivalis*’ major fimbriae interacting with β1 integrins [16]. Once inside GECs, *P. gingivalis* can rapidly localize in the perinuclear space without being constrained by membrane-bound vacuoles, where the microorganism swiftly starts associating with the endoplasmic reticulum (ER) structures to form special autophagic vacuoles which act as protective subcellular bacterial replicative niches, permitting the bacteria to gain foothold within the epithelial cells and promote subsequent intercellular spreading [15,17,18]. These host-adaptive abilities render *P. gingivalis* recalcitrant to conventional physical, chemical, or antibiotic-based control procedures [19]. Thus, specific molecular investigations need to be undertaken to effectively target intracellular *P. gingivalis*, curtail its chronic colonization of the oral cavity, and the pathologies of *P. gingivalis-*associated with systemic diseases.

Recent findings from our lab underscore a major intracellular survival mechanism where *P. gingivalis* evades ubiquitin receptor (i.e., NDP52 and P62) marking and is rapidly trafficked to the (ER) region, where its survival becomes intrinsically tied to the critical homeostatic processes of autophagy [15]. Autophagy, or the homeostatic cellular recycling mechanisms, has numerous canonical variants including microautophagy (where, typically during starvation, cellular components are directly taken up by lytic vesicles for degradation) and selective macroautophagy (where damaged cellular components or foreign bodies are ensconced by double-membraned autophagic structures called “autophagosomes” prior to being trafficked to the lysosome) that occur simultaneously in host cells [20–23]. During canonical macroautophagy, hereby dubbed “autophagy”, Beclin 1 is activated so that it can form the Beclin 1 Core Complex (comprised of Beclin 1, p150, and Vps34), which then can join with ATG14 (also known as Beclin 1-associated autophagy-related key regulator (Barkor) or ATG14L) to induce the formation of the ATG14-Involved Initiation Complex [24–28]. ATG14 has multiple N-terminal cysteine repeats that are required for the recruitment and localization of the Beclin 1 Core complex to the ER [29,30]. This complex then induces phagophore membrane formation, which is comprised of ER components and phosphatidylethanolamine (PE)-anchored, lipidated Microtubule-associated proteins 1A/1B light chain 3 (LC3) structural molecules [29,31]. LC3, a critical autophagosomal structural molecule, is traditionally primed by the protease ATG4 prior to its conjugation with PE, after which it is incorporated into the forming phagosomal membrane and lipidated [31]. These phagophores then fully form and become autophagosomes, containing the ensconced cellular components [25]. These autophagosomes then undergo maturation modification where ATG14 and other Beclin 1 Core Complex actors are removed; the autophagosomes are then are trafficked to the lysosome where ATG4 cleaves LC3, disassociating it from the autophagosomal membrane complex; and the autophagosomes are degraded following autolysosomal fusion [20,21,26,28,32–34]. If canonical selective autophagic processes specifically target invading pathogens, they are classified as xenophagy [35]. This selective variant of autophagy can be characterized by an isomer of LC3 called LC3C. Previous studies have shown that LC3C is required for the autophagic clearance of certain bacteria, such as *S. Typhimurium* [36,37], where the bacteria are targeted via ubiquitin-marking, become ensconced in LC3C-characterized autophagosomes, and are trafficked to the lysosome for degradation. However, select pathogens appear to be capable of evading xenophagy [38,39]. This evasion is not well defined but prior studies have shown that some bacteria, such as *Shigella flexneri, Vibrio parahaemolyticus, Salmonella* species, *Ralstonia solanacearum,* and *Mycobacterium species* secrete competitive or noncompetitive effector molecules, or otherwise directly inhibit canonical selective xenophagic events, such as autolysosomal fusion [39,40]. Thus, these bacteria can induce a “pro-bacterial” non-canonical variant of autophagy, where they survive within the autophagic structures*. P. gingivalis,* likewise, appears to subvert canonical selective autophagy, as recently determined by the high-resolution three-dimensional (3D) transmission electron microscopy (TEM) [15]. The initial characterizations of *P. gingivalis-* specific autophagosomes revealed that they are rich in both ER and LC3-specific autophagosomal structural molecules and contain many intact *P. gingivalis* [15]. Interestingly, *P. gingivalis* contained within these autophagosomes tend to forgo ubiquitin-marking, are not readily targeted by ubiquitin-binding-adaptor proteins such as NDP52 or P62 and are not trafficked to lysosomal degradation [15]. The high presence of LC3 in the autophagosomes could denote LC3C’s heightened involvement in this selective, non-canonical autophagy and, for the first time, suggest that LC3C has a unique role in how *P. gingivalis* usurps conventionally host-protective, antibacterial xenophagic mechanisms to prolong its survival in GECs.

Concurrently, intracellular *P. gingivalis* has been shown to promote prolonged bacterial survival via mitigating host danger signal, extracellular ATP (eATP)-induced reactive oxygen species (ROS) and the associated oxidative stress [41–44]. One of the mechanisms that intracellular *P. gingivalis* has been shown to employ is the secretion of an effector enzyme called nucleoside-diphosphate-kinase (Ndk) to inhibit eATP/P2X_7_-receptor mediated cell death in GECs via eATP hydrolysis [41,43,45]. Additionally, *P. gingivalis* can inhibit NADPH Oxidase 2-derived hydrogen peroxide (H_2_O_2_) production and subsequent antibacterial hypochlorous acid (HOCl) production [41–43]. The microorganism is able to abrogate eATP-induced ROS effects by inducing glutathione (GSH) synthesis via temporally promoting the expression of the GSH synthesis proteins glutamate cysteine ligase (GCL) subunits, GCLc and GCLm, and glutathione synthetase (GS) expression [42]. *P. gingivalis* also has been shown to inhibit apoptosis in infected GECs via activating the host anti-stress molecule heat shock protein 27 (HSp27) [46]. HSp27 is a multi-dimensional regulator protein involved in the regulation of many anti-stress pathways; HSp27 promotes actin remodeling, inhibits apoptosis, and has been shown to decrease ROS levels via reducing intracellular iron and increasing GSH levels in both HT-29 cells and CCL39 fibroblasts [47,48]. Our lab previously revealed the specific anti-apoptotic effects of HSp27 in the context of infection, as the Ndk effector of *P. gingivalis* can directly activate HSp27 via phosphorylation on its Serine (S)78 and (S)82residues to promote the survival of infected GECs [46]. Critically for this paper, however, HSp27 has recently been proposed to be involved in host protective autophagic mechanisms and activated HSp27’s induction of GSH production closely mirrors that seen during *P. gingivalis* infection in GECs [49].

Thus, in this study, we aimed to address an important knowledge gap regarding 1) the specific roles HSp27 and LC3C play the initiation of *P. gingivalis*-mediated selective autophagy in GECs, and 2) how the HSp27/LC3C coupling influences over *P. gingivalis-*mediated autophagy intertwines with the redox state of GECs to induce a pro-bacterial environment in evasion of lysosomal trafficking. Critically, we elucidate a very novel host-microbe mechanism on how *P. gingivalis* largely induces activated HSp27, which then preferentially interacts with LC3C that also becomes markedly accumulated and lipidated during *P. gingivalis* infection. The temporal phosphorylation of HSp27 (P-HSp27) by the bacterial Ndk later points in the infection leads to strengthened partnering between P-HSp27 and LC3C protects LC3C’s C-terminal tail from canonical protease cleavage and promotes the highly extended tethering of the ATG14-Involved Initiation Complex to *P. gingivalis-*specific autophagosomes. Additionally, we demonstrate that these pro-bacterial autophagic associations are critically tied to the host cell’s redox state, as HSp27 jointly acts as a redox homeostatic regulator to mitigate the oxidative stress caused by anti-bacterial host responses such as eATP. These findings elucidate potential avenues for developing effective therapeutic strategies to control the chronic colonization of *P. gingivalis* in the oral cavity and identify critically novel autophagic interactors that could be jointly therapeutically targeted to curtail the other pathogenic bacteria that usurp autophagy to cause disease.

## RESULTS

### *P. gingivalis* infection readily influences the levels and localization patterns of the anti-stress host molecule HSp27

Previous studies have shown that *P. gingivalis* can activate HSp27 to promote bacterial survival via inhibiting apoptosis of infected GECs [46]. Thus, initial western blotting and confocal microscopy were performed to define how *P. gingivalis* infection specifically influences the levels and presence of HSp27 in GECs. *P. gingivalis* infection induced a large accumulation and spatial reorganization of HSp27 in GECs, as HSp27 shifted from being diffused throughout the cytoplasm to being highly localized with intracellular *P. gingivalis* in the ER region (**Fig. 1A and 1B**). The ready co-localization (Pearson, r>0.87 for infected GECs; Two-Tailed Student T-Test, p<0.05) suggests a potential pro-bacterial interaction between HSp27 and ER-centric, P*. gingivalis*-specific autophagosomes (**Fig. 1A and 1Ai**).

**Figure 1.**
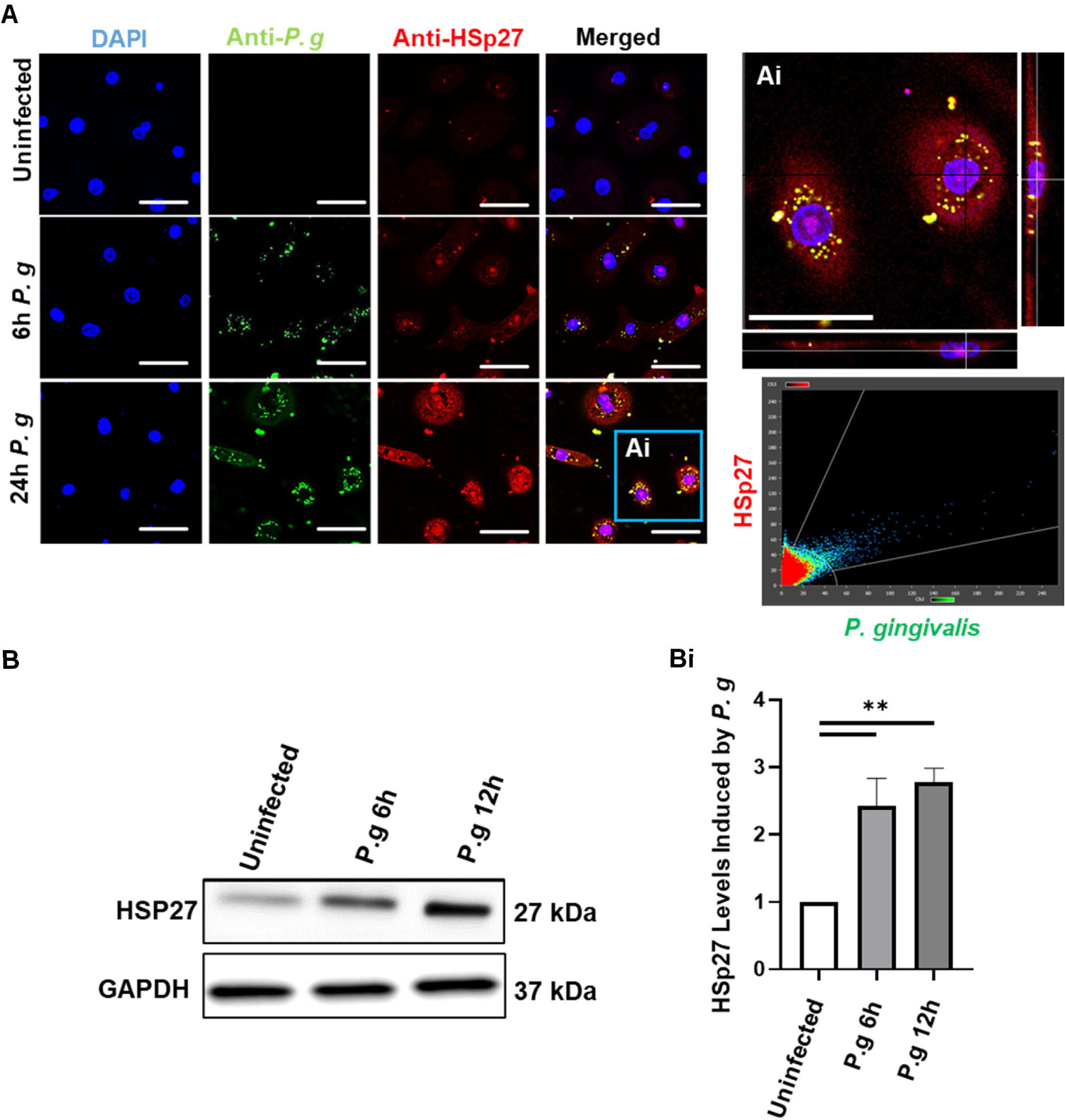
*P. gingivalis (P. g)* Infection Causes a Significant Increase in the Protein HSp27, Accompanied by Large Spatial Accumulation of Hsp27 with the Bacteria in a Temporal Manner in Primary GECs. (**A**) Representative confocal microscopy images of *P. g*-infected human primary GECs at an MOI 100, at 6 h and 24 h after infection. GECs were then stained for *P. g* (rabbit anti-P. g; Alexa 488; green) or HSp27 (mouse anti-HSp27; Alexa 568; red). GECs were then imaged via the Leica DM6 CS Stellaris 5 Confocal/Multiphoton System at 63x. (**Ai**) Imaris Software was used to create a zoomed image of infected GECs and was used to calculate the amount of co-localization between *P. g* and HSp27. HSp27 was found to readily colocalize with *P. g*, having an average Pearson correlation coefficient of 0.87 via the Imaris software. Scale bar is 30 µm for 63x and Zoomed Magnification. **(B**) *P. g* was added at MOI 100 to GECs, which were incubated 6 or 12h. Cell lysates were then analyzed via western blot. (**Bi**) Quantitative ImageJ analyses of western blot results. Data is represented as Mean±SD, where n=3 and p<0.05 was considered as significant via Two-Tailed Student T-test. *p<0.05.

### HSp27 presence is vital to *P. gingivalis-*mediated non-canonical (pro-bacterial) autophagy

Given the apparent influence of *P. gingivalis* infection on HSp27 levels (**Fig. 1**) and HSp27’s suggested involvement in host protective autophagic mechanisms [49], it became prudent to determine how HSp27 depletion would affect *P. gingivalis*-induced non-canonical autophagy. To do so, a novel process was developed to specifically elucidate HSp27’s relevance, where only *P. gingivalis-*specific autophagosomes were selectively isolated from either native GECs or GECs depleted of HSp27 via targeted siRNA (Methodology and Proof of Principle shown in **Fig. 2A and 2B**) [50]. Prior to experimental implementation, precautions were taken to ensure that the lipobiotin treatments and magnetic labelling of the isolation processes did not interfere with the invasive capability or survival of *P. gingivalis* (**Fig. S1**). The isolated intact *P. gingivalis*-specific autophagosome findings were confirmed via imaging and comparing HSp27-depleted, infected GECs to non-depleted infected GECs with high-resolution confocal microscopy.

**Figure 2.**
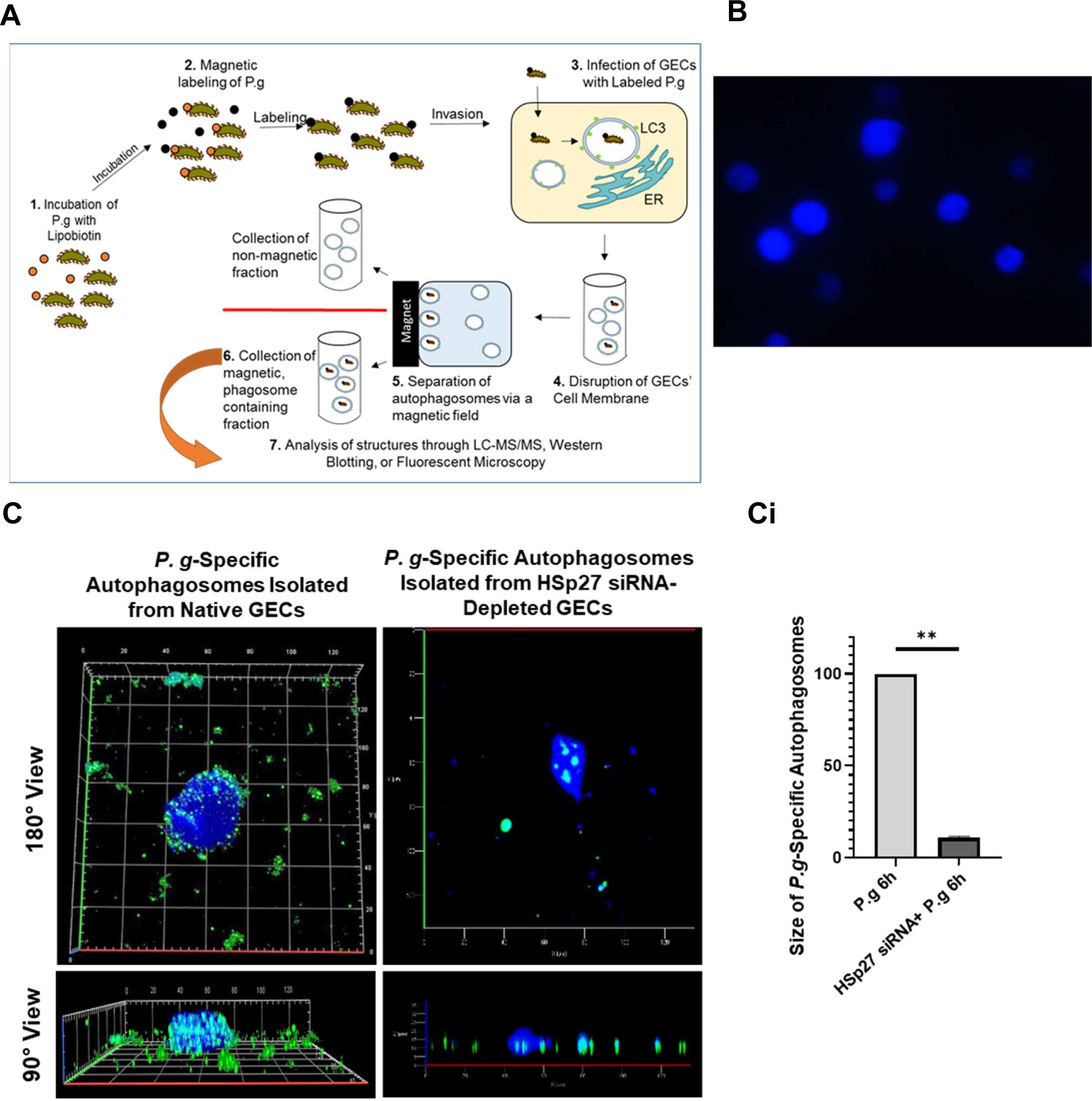
The Integrity of *P. gingivalis (P. g)*-Specific Autophagosomes is Highly Dependent on HSp27 Presence. Human primary GECs were treated with HSP27siRNA (100nM) for 48 h prior to incubation with *P. g (*MOI 100) for 6 h. Autophagosomes were then isolated and analyzed via Confocal Microscopy. (**A**) Schematic autophagosomal isolation method of infected GECs. (**B**) Confocal microscopy images of autophagosomes (ThiolTracker Violet; blue) were obtained via Super Resolution Zeiss Airyscan LSM 880 at 20x. (**C**) Autophagosomes were stained for *P. g* (rabbit anti-*P. g*; Alexa 488; green) and reduced GSH (ThiolTracker Violet; blue). Confocal microscopy images of autophagosomes were obtained via Super Resolution Zeiss Airyscan LSM 880 at 20x. (**Ci**) Quantitative ImageJ analysis of Confocal microscopy results was then performed. Data is represented as Mean±SD, where n=25 and p<0.05 was considered as statistically significant (Student two-tailed T-test). **p<.005

Initial confocal image analyses showed that there were ubiquitous deformations in the circular shape of intact *P. gingivalis*-specific autophagosomes following HSp27 depletion by targeted siRNA (**Fig. 2C**) and an 8-fold decrease in average autophagosomal size, when compared to non-depleted counterparts (**Fig. 2Ci**). ThiolTracker, a probe for intracellular thiols and glutathione (GSH) levels, was utilized to examine the intact morphology of *P. gingivalis*-specific autophagosomes when compared to their HSp27-depleted counterparts, visualize their GSH-rich nature, and highlight the difference of size and shape between the two treatment groups (**Fig. 2B and 2C**). Provided these findings, transmission electron microscopy (TEM) was utilized to confirm that, following HSp27 depletion, *P. gingivalis*-specific autophagosomes underwent canonical, anti-bacterial lysosomal trafficking (**Fig. 3A**), thus confirming HSp27’s relevance in *P. gingivalis-*induced autophagy.

**Figure 3.**
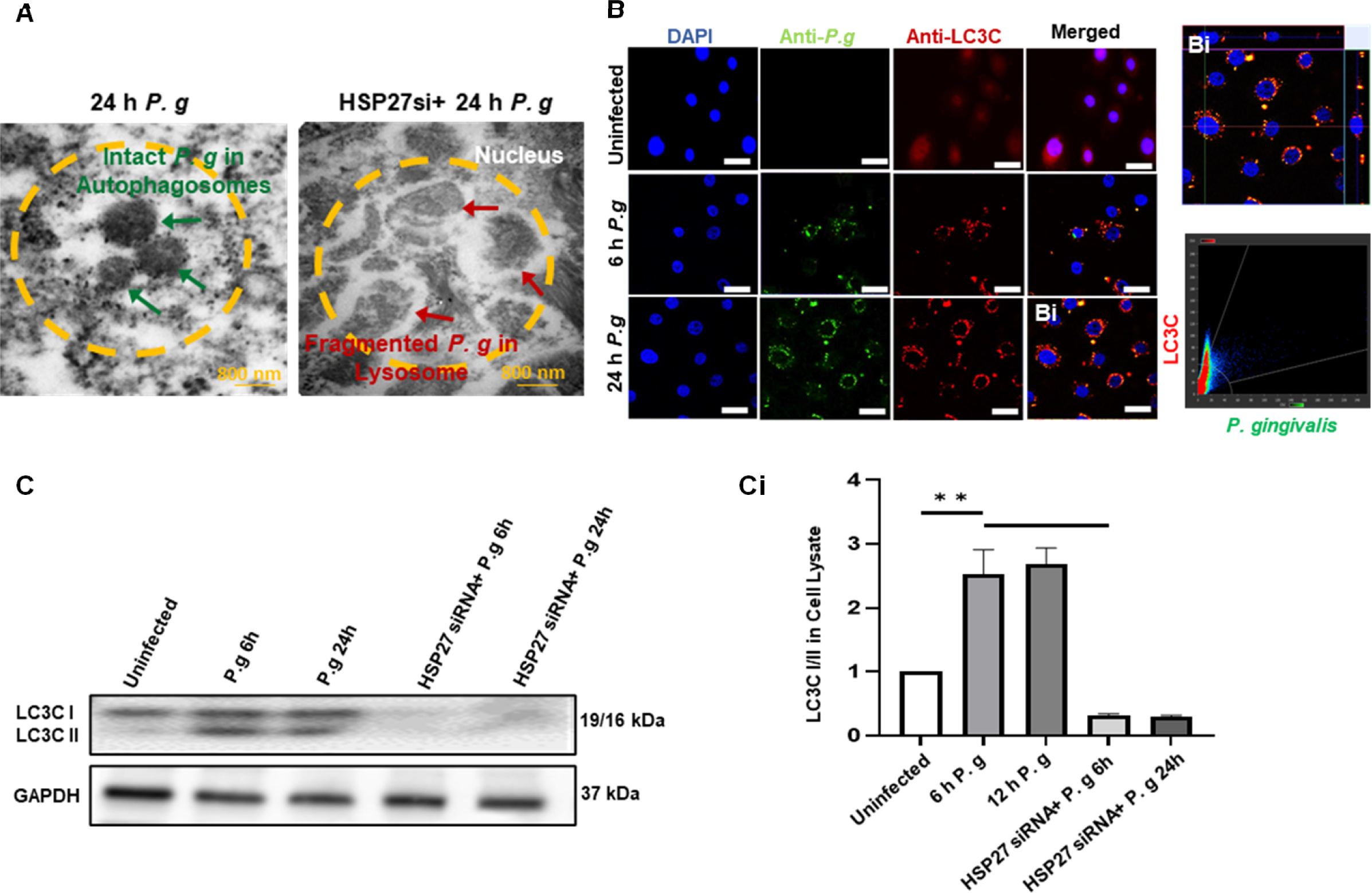
Intracellular *P. gingivalis* (*P. g*) Significantly Induces and Co-Localizes with LC3C, an Isomer of LC3, and this Specific Event is Highly Dependent on HSp27 for Successful Autophagic Survival. Human primary GECs were treated with HSp27 siRNA (100nM) for 48 h. *P. g* was added at MOI 100 to GECs, which were incubated for 6 or 24 h. (**A**) GECs were targeted for *P. g* (rabbit anti-*P. g*; goat anti-rabbit Ultra Small Gold Antibody) and labeled *P. g* was found to be readily ensconced within double-membraned autophagosomes in GECs. Following HSp27 depletion, *P. g* appeared to readily start to degrade. Representative transmission electron microscopy images of *P. g*-infected GECs were also taken at 80 kV and 100000x magnification. Scale bar is 800 nm. (**B**) 6 h and 24h *P.* g-infected GECs were also stained for *P. g* (rabbit anti-P*. g*; Alexa 488; green) and LC3C (mouse anti-LC3C; Alexa 568; red) to examine whether LC3C characterizes *P. g*-specific autophagosomes. These cells were then imaged via confocal microscopy (Leica DM6 CS Stellaris 5 Confocal/Multiphoton System) at 63x. The range of z-stacks was kept consistent and representative images were selected from the mid-ranged sections. (**Bi**) The Imaris software was utilized to obtain a xoomed 63x Orthogonal Image of 24 h *P. g* infection and found heightened co-localization between *P. g* and LC3C. LC3C was found to readily colocalize with *P. g,* having an average Pearson correlation coefficient of 0.96 via the Imaris post-processing software. (**C**) Lysates of infected and HSp27-depleted GECs were also analyzed via western blotting. Non-target controls were performed and not shown. (Ci) Quantitative ImageJ analysis was performed for the western blot results. Data is represented as Mean±SD, where n=3 and p<0.05 was considered as statistically significant via Student two-tailed T-test. **p<.005.

### *P. gingivalis-*mediated non-canonical autophagy is characterized by LC3C, a novel isoform of the autophagosome structural molecule LC3C

Prior studies have recently associated the LC3C isoform of the autophagosomal structural molecule, LC3, to xenophagic mechanisms [36], thus, we wished to determine what role LC3C might play in *P. gingivalis-*induced, non-canonical autophagy. Confocal microscopy was used to visualized how LC3C spatiotemporal localization were affected by *P. gingivalis* infection in GECs: LC3C presence was significantly increased in infected GECs and differed in subcellular localization patterns when compared to uninfected counterparts (**Fig. 3B and 3Bi**). *P. gingivalis* infection induced a shift in LC3C from the nuclear region to the perinuclear region, where it highly co-localized with intracellular *P. gingivalis* localized to the ER region (Pearson, r>0.96 for infected GECs; Two-Tailed Student T-Test, p<0.05). Western blotting results showed that LC3C levels (LC3C I) and LC3C lipidation (LC3C II), a signifier of LC3C-specific autophagosomal formation and flux [36,37], were readily increased upon *P. gingivalis* infection (**Fig. 3C and 3Ci**). Interestingly, LC3A/B II levels were not majorly impacted *P. gingivalis* infection (**Fig. S2A**), LC3B depletion via siRNA did not influence intracellular *P. gingivalis* survival (**Fig. S3B**), LC3C levels and lipidation were only increased upon the infection while there was not any detectable induction in the LC3C levels and lipidation following the GECs being stressed via starvation (**Fig. S3C1**). These combined findings suggest that the induction in LC3C is solely driven by *P. gingivalis-*induced non-canonical autophagy, and that *P. gingivalis*-induced autophagy in GECs is not majorly reliant on the other LC3 isoforms.

### The non-canonical, *P. gingivalis-*specific, induction of LC3C is dependent upon the presence of HSp27

Next, we wished to determine how HSp27 depletion impacts LC3C-characterized, *P. gingivalis*-induced autophagy. HSp27 depletion via targeted siRNA, resulted in the decrease of LC3C I/II, signifying that LC3C-rich autophagosomes were either no longer readily forming or were being degraded (**Fig. 3C and 3Ci**). To reconfirm these theorized trafficking patterns of *P. gingivalis-*specific, LC3C-rich autophagosomes following HSp27 depletion, a novel construct (mCherry-GFP-LC3C, where LC3C was conjugated to an acid sensitive eGFP and an acid insensitive mCherry probe) was created to permit the visualization of autolysosomal fusion and resulting lysosomal acidification [51]. If the construct-comprised, *P. gingivalis*-specific autophagosomes fuse with the lysosome, the acid sensitive green fluorescence would quench, resulting in only the red fluorescence being visible. *P. gingivalis* was found to readily co-localize with the LC3C construct (**Fig. 4A**; Pearson, r>0.82 for infected GECs; Two-Tailed Student T-Test, p<0.05). Additionally, *P. gingivalis*-specific autophagosomes did not appear undergo canonical lysosomal acidification, as both green and red probes substantially fluoresced in infected GECs (**Fig. 4A**). However, following HSp27 depletion, the green fluorescence quenched, suggesting that canonical, antibacterial autophagy was prevailing and that the autophagic *P. gingivalis* was undergoing lysosomal degradation (**Fig. 4A**). To supplement these findings, traditional immunofluorescence staining for *P. gingivalis* and the lysosomal membrane protein LAMP-1 was performed. These results showed that *P. gingivalis* only associated to lysosomal compartments following HSp27 depletion via targeted siRNA (**Fig. 4B**; Pearson, r<0.25 for infected GECs versus r>0.83 for HSp27-depleted, infected GECs; Two-Tailed Student T-Test, p<0.05). To confirm the non-canonical, pro-bacterial autophagic trafficking patterns seen following *P. gingivalis* infection, autophagic inhibitors (bafilomycin A1, an inhibitor of vacuolar H+-ATPase that blocks lysosome fusion [52]; 3-Methyladenine (3-MA), a selective inhibitor for the PI3K protiensVps34 and PI3Kγ [53]; and pepstatin A methyl ester, a cathepsin D/lysosomal protease inhibitor [54]) were individually utilized in conjunction with HSp27 depletion. TEM analysis, LAMP-1 immunofluorescence staining, and the LC3C reporter system were used to examine the effects of the autophagic inhibitors on *P. gingivalis-*specific autophagic trafficking. Treatment of HSp27-depleted, infected GECs with bafilomycin A1 and pepstatin A methyl ester--late-stage autophagic inhibitors--resulted similar visual trafficking patterns to non-HSp27-depleted, infected GECs, where GECs exhibited no quenching of the acid-sensitive eGFP probe in confocal microscopy analyses, did not localize to LAMP-1, and did not undergo lysosomal degradation in TEM analyses (**Fig. 4A-4C**). These results signify that *P. gingivalis-*specific autophagosomes do not appear to canonically mature nor undergo canonical lysosomal fusion in GECs.

**Figure 4.**
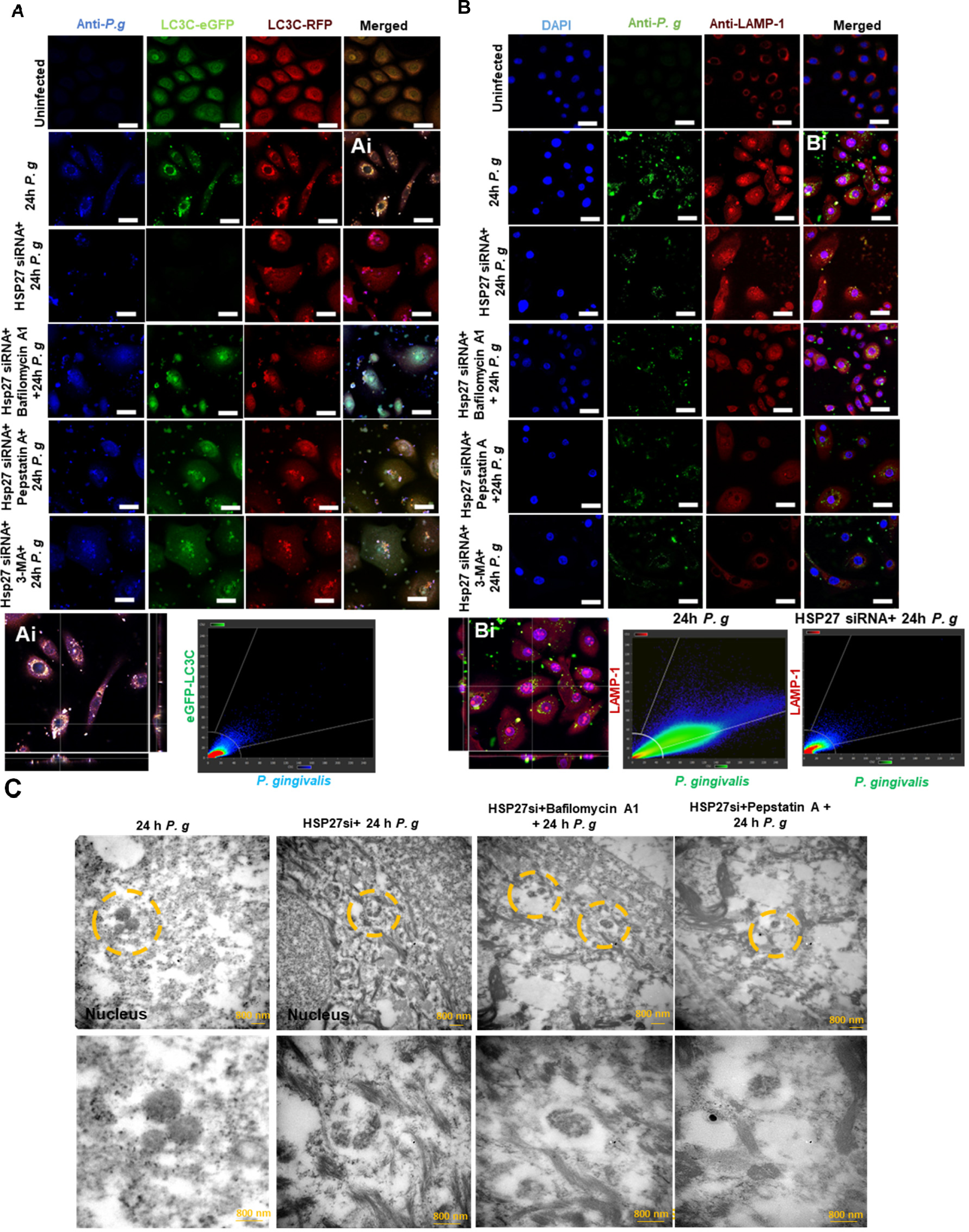
HSp27 Presence Permits the Prolonged Existence of LC3C-characterized, *P. gingivalis* Specific Autophagosomes by Hampering Canonical Autolysosomal Fusion in Primary GECs. (**A**) Human primary GECs were transfected with mCherry-eGFP-LC3C for 48 h. Select GECs were also treated with 1 uM of the autophagolysosomal fusion inhibitor Bafilomycin A1, 1 uM Pepstatin A, or 5 mM 3-MA. Others were treated with Hsp27 siRNA (100nM) for 24 h. *P. g* was added at MOI 100 to GECs, which were incubated for 24 h. (**A**) GECs were then stained for *P. g* (mouse anti-*P.g;* Alexa 405; blue) and were mounted. GECs were then imaged via confocal microscopy (Leica DM6 CS Stellaris 5 Confocal/Multiphoton System) at 63x. (**Ai**) Imaris was used to obtain a zoomed 63x orthogonal image of 24 h *P. g* infection and measure the high co-localization levels between *P. g* and the LC3C Reporter System. *P. g* localized readily to the LC3C construct, with a Pearson correlation coefficient of 0.82. **(B)** Separately, GECs were additionally stained for *P. g* (rabbit anti-P*. gingivalis*; Alexa 488; green) and LAMP-1 (mouse anti-LAMP-1; Alexa 568; red) and were imaged. The range of all z-stacks was kept consistent and representative images were selected from the mid-ranged sections. Scale bar is 40 µm for 63x Magnification. (**Bi**) Imaris was used to obtain a zoomed 63x orthogonal image of 24 h *P. g* infection and measure the co-localization levels between *P. g* and LAMP-1 in infected and treated GECs. While *P. g* infected GECs did not exhibit high co-localization with LAMP-1 (Pearson correlation coefficient of .25), their HSp27-depleted counterparts did, with an average Pearson correlation coefficient of 0.83. The Scale bar is 20 µm for 63x Magnification. (**C**) Finally, GECs treated with autophagic inhibitors were targeted for *P. g* (rabbit anti-*P. g*; goat anti-rabbit Ultra Small Gold Antibody) and labeled *P. g* was found to be readily ensconced within double-membraned autophagosomes in GECs. Representative transmission electron microscopy images of *P. g*-infected GECs were taken at 80 kV and 30000x or 100000x magnification. Following HSp27 depletion, *P. g* appeared to readily start to degrade, however treatment with late-stage autophagic inhibitors Bafilomycin A1 or Pepstatin A appeared to rescue *P. g* from degradation. Representative transmission electron microscopy images of *P. g*-infected GECs were taken at 80 kV and 30000x or 100000x magnification. Scale bar is 800 nm.

### *P. gingivalis-*mediated non-canonical autophagy is uniquely dependent upon the presence of the LC3C isoform

Due to the very specific induction of LC3C caused by *P. gingivalis* and the high levels of co-localization with intracellular *P. gingivalis* and LC3C seen GECs, we wished to reconfirm LC3C’s relevance explicitly for the autophagic survival of *P. gingivalis*. To do so, *P. gingivalis-* specific autophagosomes were isolated from LC3C-depleted, infected GECs, so that they could be compared to non-depleted counterparts. Additionally, an *in situ* antibiotic protection assay using 16S rRNA probes specific to *P. gingivalis* was used to measure the levels of live intracellular *P. gingivalis* survival in LC3C-depleted, infected GECs [55]. Results showed that following LC3C depletion via targeted siRNA, the integrity of *P. gingivalis-*induced non-canonical autophagy collapsed: no intact *P. gingivalis* specific autophagosomes could be readily isolated (**Fig. 5A**). Additionally, the levels of live intracellular *P. gingivalis* were significantly decreased following LC3C depletion (**Fig. 5B**). These results combined with those confirming HSp27’s necessity in *P. gingivalis* induced autophagy suggest that LC3C is 1) a key structural component required for the viability of *P. gingivalis-*driven non-canonical autophagy and 2) that HSp27 and LC3C are mutually inclusive and essential for *P. gingivalis*’ autophagic survival. The removal of either of these key players results in the disintegration of the autophagic integrity of *P. gingivalis*.

**Figure 5.**
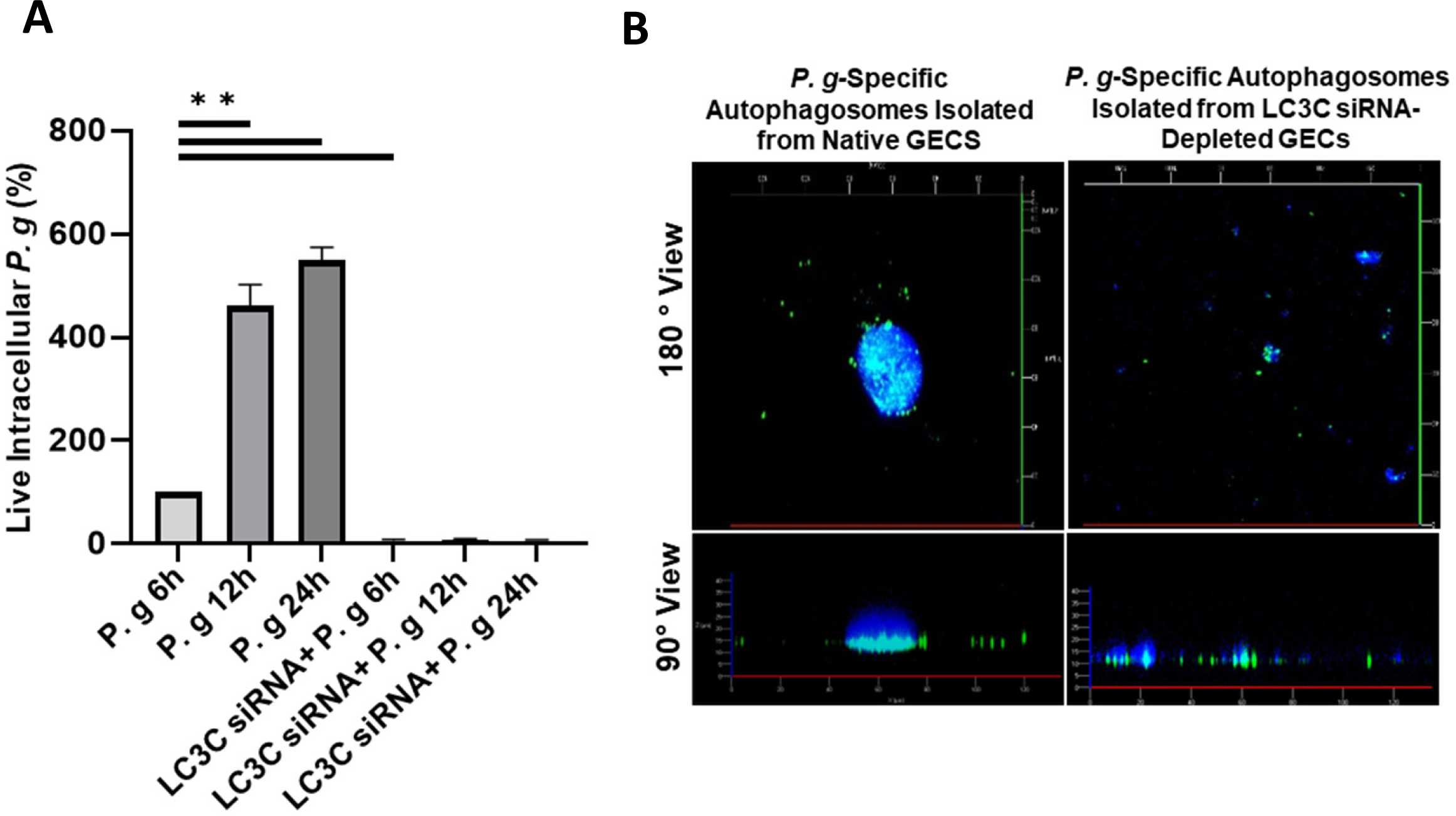
Depletion of LC3C via siRNA Collapses *P. gingivalis (P. g)*-Induced Non-Canonical Autophagosomal Integrity. Human primary GECs were treated with LC3C siRNA (100nM) for 48h. *P. g* was added at MOI 100 to GECs for 6, 12, or 24 h. (**A**) Intracellular *P. g* survival after LC3C siRNA depletion was determined using a standard antibiotic protection assay. In brief, any extracellular bacteria were killed via 1h gentamicin (300 μg/mL) and metronidazole (200 μg/mL) treatment. cDNAs were synthesized for qPCR using *P. g*-specific 16S rRNA primers to quantify intracellular levels of live *P. g*. Data is represented as Mean±SD, where n=3 and p<0.05 was considered as statistically significant via Student two-tailed T-test. *p<.05 **p<.005. (**B**) *P. g*-specific autophagosomes were also selectively isolated. Autophagosomes were stained for *P. g* (rabbit anti-*P. g*; Alexa 488; green) and reduced GSH (ThiolTracker Violet; blue). Confocal images of *P. g*-specific autophagosomes at 6 h post-infection (63x) were taken utilizing the Super Resolution Zeiss Airyscan LSM 880.

### HSp27 interacts with LC3C in *P. gingivalis-*induced autophagy in a GSH-mediated, redox dependent mode

Given *P. gingivalis’* known capability to abrogate host cell redox stress and HSp27’s established status as an anti-stress modulator, the potential role of HSp27 in the inhibition of oxidative stress during *P. gingivalis-*induced, non-canonical autophagy needed to be separately elucidated. Extracellular ATP (or eATP), a well-established, physiologically-relevant ROS-inducer abrogated by *P. gingivalis* in human primary GECs [42,45,46,55,56], was utilized here to define HSp27’s theoretical role as a redox regulator as well as an autophagic influencer. Western blotting was performed, and the findings showed that the *P. gingivalis*-infected GECs substantially maintained the induced LC3C I/II levels upon eATP treatments, while decreased levels of LC3C I/II following Hsp27 depletion were further synergized by treatment with eATP in GECs (**Fig. 6**). This heightened reduction underpins the dual role HSp27 appears to play in *P. gingivalis*-induced non-canonical autophagy and host-pathobiont symbiosis; HSp27 could be both influencing the formation of LC3C-rich autophagosomes while also acting as a redox stabilizing molecule in infected GECs.

**Figure 6.**
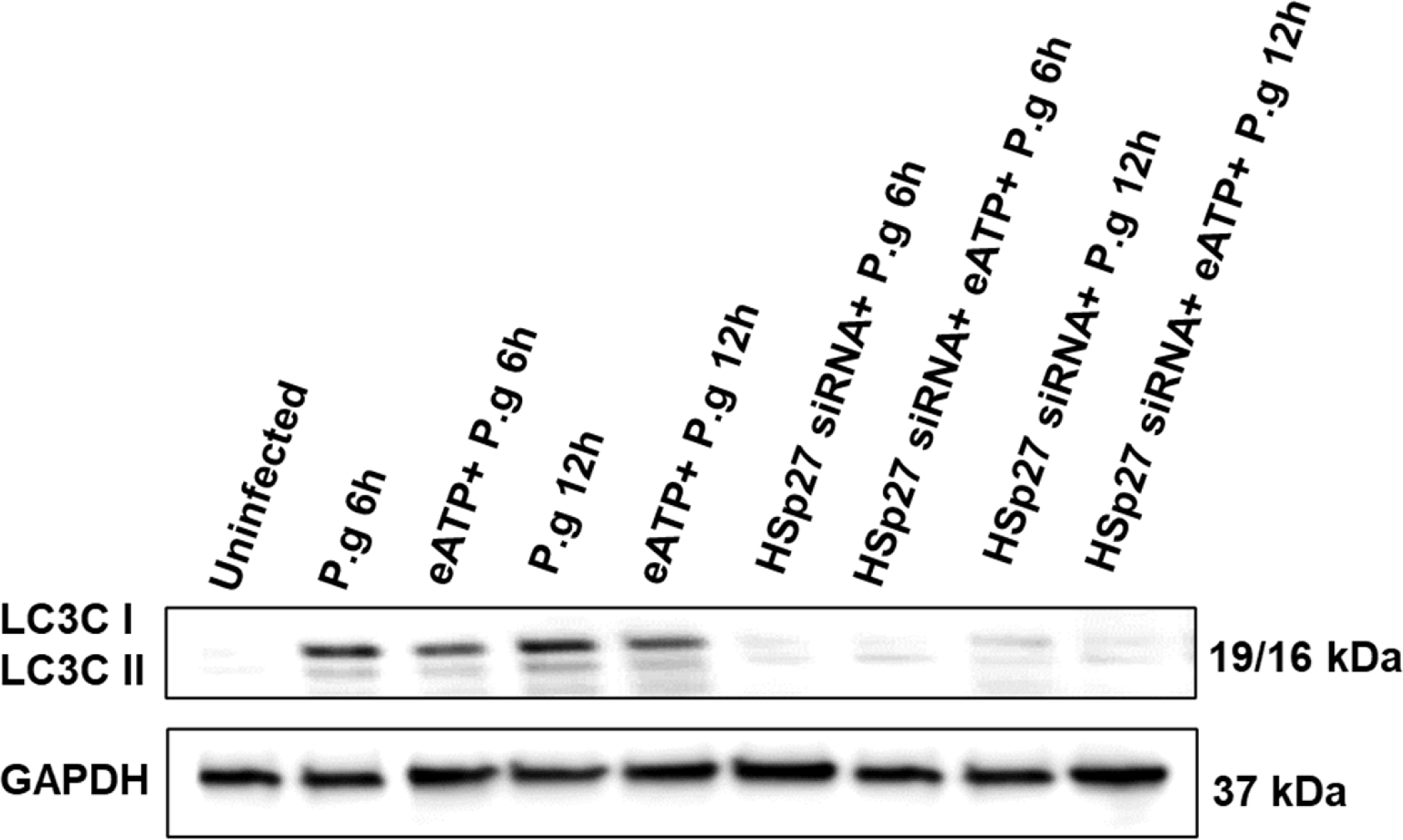
*P. gingivalis (P. g)* Causes the Nucleation of Hsp27-Mediated LC3C Accumulation and Lipidation; this Specific Assembly is Highly Dependent on Host Cells’ Redox Potential Determined by eATP Treatments. Human primary GECs were treated with HSp27 siRNA (100nM) for 48 h. *P. g* was added at MOI 100 to GECs, which were incubated 6 and 24 h. Non-Depleted and HSp27-depleted GECs also were treated with the physiologically-relevant oxidative stress inducer eATP (3mM) treatment for 30 min prior to infection, and were analyzed by western blot.

Previous studies note that HSp27 can act as a redox stabilizer via inducing GSH levels [47,57]. As *P. gingivalis* can also temporally influence GSH synthesis to abrogate anti-bacterial HOCl production [42] and our intact *P. gingivalis-*specific autophagosomes were shown to be enriched in GSH, when compared to HSp27-depleted counterparts (**Fig. 2B and 2C**), we measured the GSH levels following *P. gingivalis* infection and HSp27 depletion to see if HSp27 is inducing GSH during *P. gingivalis* infection in GECs. GSH was found to be induced by *P. gingivalis* in a manner that was dependent upon HSp27 presence as, following HSp27 depletion, GSH presence fell below the levels found in untreated, uninfected GECs (**Fig. 7A**). Given GSH’s apparent relevance, we then wanted to find an alternate means to induce GSH synthesis independent of HSp27 presence, to see if HSp27’s effects on GSH could be restored. N-acetyl Cysteine (NAC), a universal antioxidant and supplier of the rate-limiting GSH precursor cysteine [58,59], was identified as a potential GSH-inducer, so GECs were treated with exogenous NAC prior to *P. gingivalis* infection (the optimum concentration of NAC used as previously determined in GECs [41]. It was found that NAC treatment promoted a mild restoration of GSH levels following HSP27 depletion (**Fig. 7A**).

**Figure 7.**
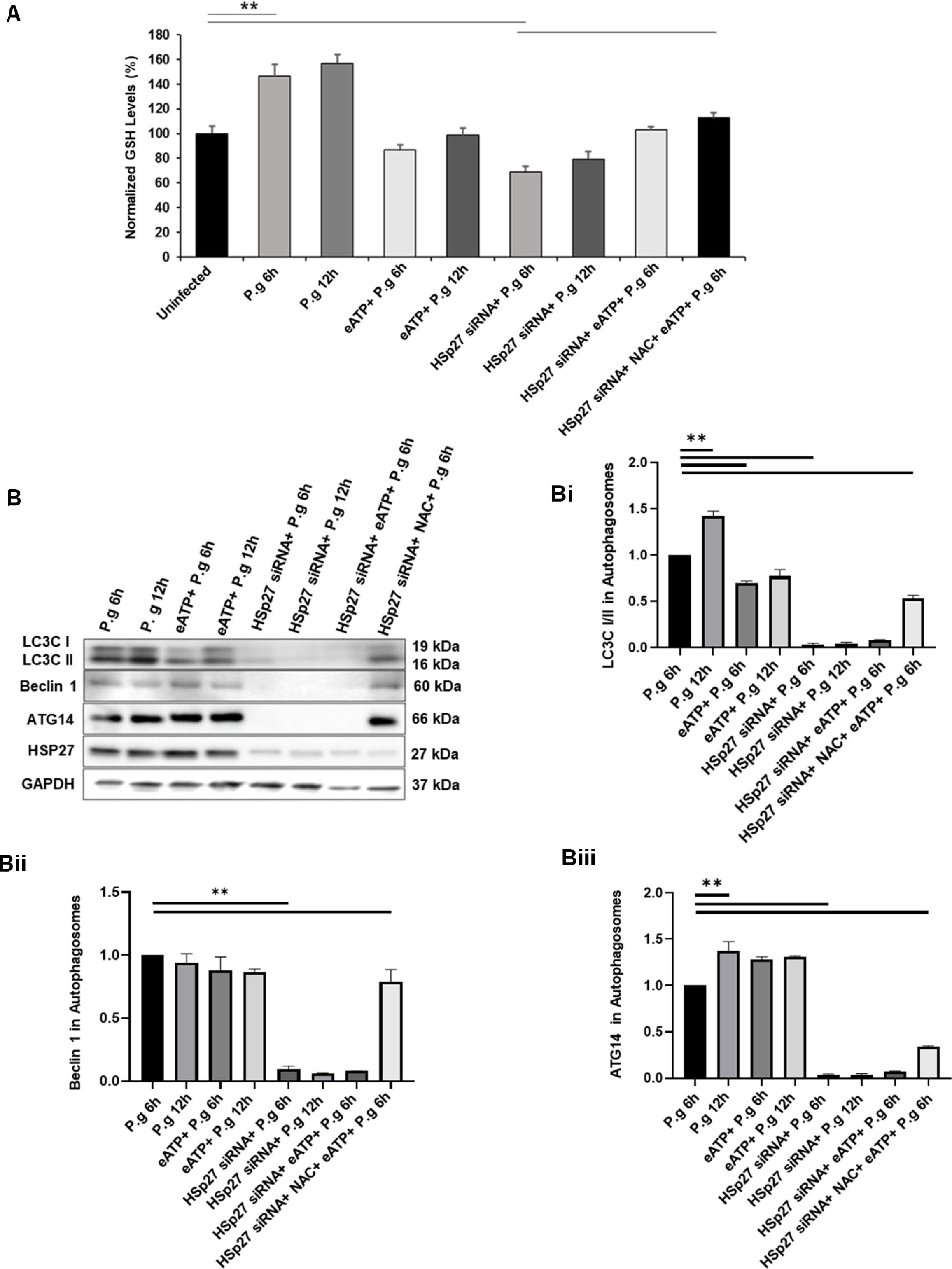
*P. gingivalis* (*P. g*) Induces and Prolongs the Autophagosomal LC3C/Beclin 1/ATG14 Nucleation Complex in a Manner Dependent upon HSp27 and the Reduced Redox State of Infected GECs as Determined by Isolated *P. g-*Specific Autophagosomes. GECs were treated with HSP27siRNA (100nM) for 48 h. Select GECs were also treated with N-acetyl Cysteine (NAC)(50 uM) for 1 h and/or eATP (3mM) for 30 min. *P. g* was added at MOI 100 to GECs, which were incubated 6 and 12 h. Autophagosomes were then isolated and prepared for analysis. (**A**) The glutathione (GSH) levels of primary GECs were also measured using chemiluminescence detection. (**B)** Isolated autophagosomes were analyzed via western blot. (**Bi**), (**Bii**), (**Biii**) Quantitative ImageJ analysis was performed of each of the western blot results. Data is represented as Mean±SD, where n=3 for results. p<0.05 was considered as statistically significant via Student two-tailed T-test. *p<.05 **p<.005

With these novel redox results accounted for, we wished to specifically examine the LC3C-HSp27 interactions in the context of redox-stress. Western blotting of isolated *P. gingivalis-*specific autophagosomes was performed. These results showed similar trends to those of the cell lysates (**Fig. 6**), where eATP treatments further decreased LC3C presence and lipidation in autophagosomes isolated from HSp27-depleted GECS (**Fig. 7B**). Importantly, however, the novel NAC treatment of HSp27-depleted, eATP-induced oxidative-stressed GECs resulted in a partial restoring of LC3C I/II levels in isolated *P. gingivalis*-specific autophagosomes (**Fig. 7B and 7Bi**). These results suggest a partial restoration of an altered pro-bacterial autophagic flux that is caused by the return of a favorable redox environment.

### HSp27 and LC3C maintain the ATG14-Involved Initiation Complex assembly during *P. gingivalis-*mediated autophagy

Further examinations were then performed to define the mechanisms of how HSp27 and LC3C might directly mediate the autophagic lifestyle of *P. gingivalis* in infected GECs. To do so, and provided GSH-mediated oxidative stress’ relevance during *P.* gingivalis-induced autophagy, *P. gingivalis-*specific autophagosomes were isolated from both native, infected GECs and GECs that were treated with the GSH-synthesis inhibitor buthionine sulfoximine (BSO); these autophagosomes were then analyzed via label-free-quantification LC-MS/MS. The total LC3 levels (the isoforms were not separately identified) and HSp27 levels were found to be enriched in *P. gingivalis* specific autophagosomes and were readily decreased in GSH-depleted, *P. gingivalis-*specific autophagosomes (**Fig. 3SA and 3SB**). Interestingly, however, both Beclin 1 and ATG14, critical ATG14-Involved Initiation Complex proteins [32,33], were also found to be highly enriched at all time points of *P. gingivalis* infection and were only diminished following BSO treatment (**Fig. 3SC and 3SD**). ATG14’s prolonged and heightened presence in the *P. gingivalis-*specific autophagosomes drew particular interest as, canonically, ATG14 is a transient modulator of the Beclin 1 Core Complex which is suggested to be sequentially removed during canonical autophagosomal maturation and exhibits a preference for binding to LC3C *in vivo* [30]. These patterns suggest that autophagic nucleation, the initial formation of autophagosomes from initial single-membrane phagophore structures to autophagosomes, was being modified by *P. gingivalis* infection and needed to be examined in more detail. Thus, isolated, *P. gingivalis-*specific autophagosomes were analyzed via western blot. Both Beclin 1 and ATG14 levels were found to be temporally increased by *P. gingivalis* infection in GECs and were notably diminished following HSp27 depletion (**Fig. 7B and 7Bii-7Biii**). Additionally, LC3C, Beclin 1 and ATG14 levels were partially restored following treatment of HSp27-depleted, eATP-stressed cells with NAC (**Fig. 7)**. These findings highlight HSp27 and LC3C’s specific relevance in inducing and maintaining a redox-sensitive autophagic nucleation complex unique to *P. gingivalis-*induced pro-bacterial autophagy induces.

### HSP27 and LC3C specifically interact with ATG14 and Beclin 1 to maintain ATG14-Involved Initiation Complex formation

Given the vital roles of HSp27 and LC3C appear to have in maintaining the ATG14-Involved Initiation Complex during *P. gingivalis-*induced autophagy in GECs (**Fig. 7**), we then wanted to mechanistically define the pro-bacterial interactions between HSp27, Beclin 1, ATG 14, and LC3C. To do so, a co-immunoprecipitation pulldown assay was performed utilizing an anti-LC3C antibody. Western-blotting showed that *P. gingivalis* infection induced the direct formation of a HSp27-LC3C-Beclin 1-ATG14 pro-bacterial complex (**Fig. 8**). Interactions between LC3C and the autophagic initiators were strictly dependent upon HSp27 presence and the host cell’s over-all redox state, as following HSp27 depletion, no interactions between the autophagic molecules were found. NAC treatment of HSp27-depleted, oxidatively stressed GECs, however, was capable of a partial rescuing of pro-bacterial partnering (**Fig. 8A**). These molecular interactions were additionally confirmed via immunofluorescence, where strong co-localizations of HSp27, Beclin 1, LC3C, and ATG14 were clearly visualized following prolonged infection with *P. gingivalis* (**Fig. 8B**; Pearson, r<0.83 for infected GECs versus r>.25 for HSp27-depleted, infected GECs; Two-Tailed Student T-Test, p<0.05).

**Figure 8.**
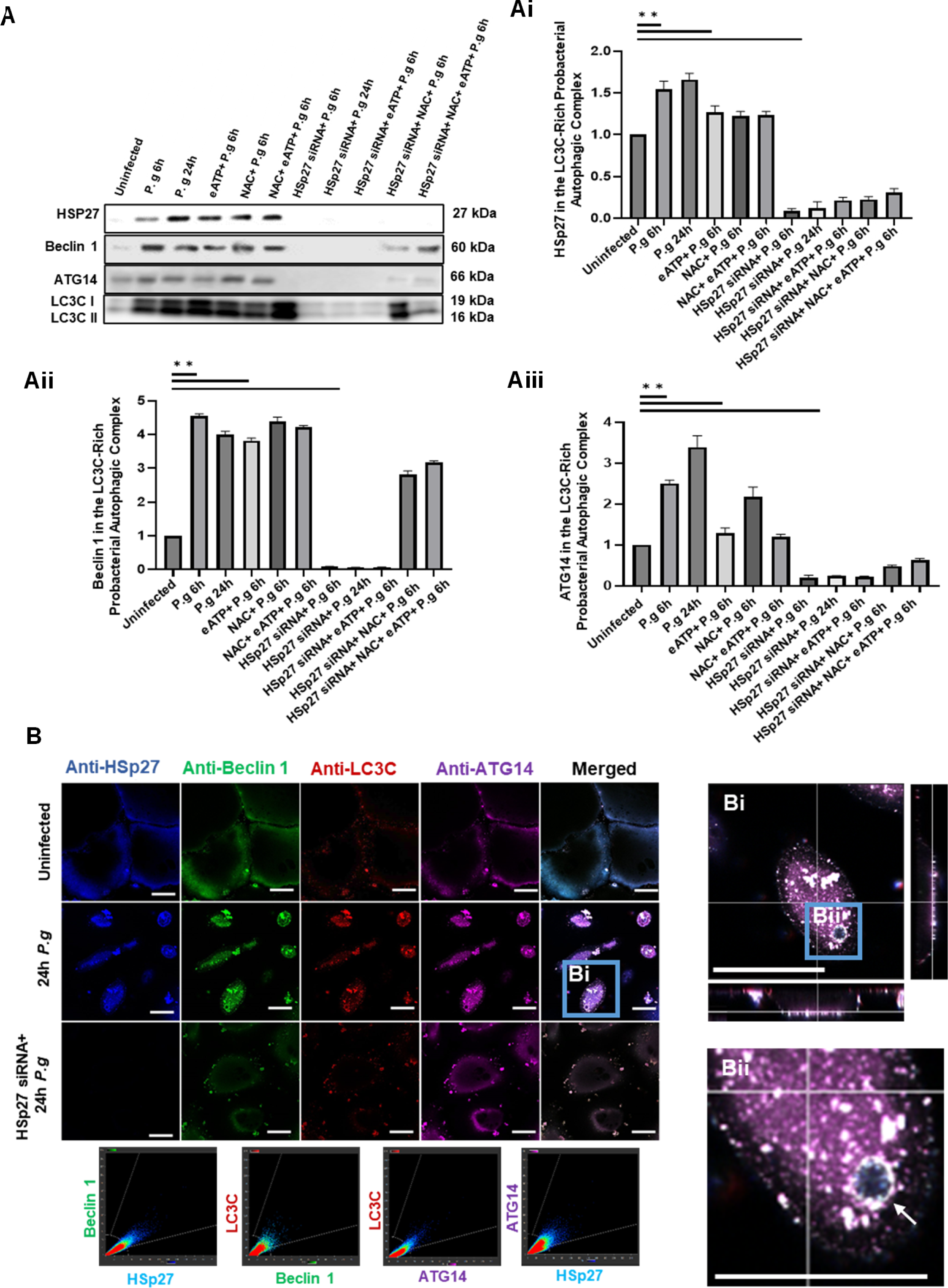
HSp27 and LC3C Recruit Beclin 1 and ATG14 to Form a Temporal Pro-bacterial Autophagic Complex, which Can Be Disrupted by Increased Oxidative Stress. Human Primary GECs were treated with HSp27siRNA (100nM) for 48 h. Select GECs were also treated with N-acetyl Cysteine (NAC) (50 uM) for 1 h and/or eATP (3mM) for 30 min. *P. gingivalis (P. g)* was added at MOI 100 to GECs, which were incubated 6 and 12 h. GECs were then lysed and the extracts were incubated in rabbit anti-LC3C antibody over-night. Samples underwent co-immunoprecipitation. (**A**) The eluted protein complexes were then analyzed by western blot. (**Ai**), (**Aii**), and (**Aiii**) Quantitative ImageJ analysis of western blot results was performed for each of the proteins in question. (**B**) GECs also underwent staining for HSp27 (goat anti-HSp27; Alexa 405; blue), LC3C (rabbit anti-LC3C; Alexa 488; green), and Beclin 1 (sheep anti-Beclin 1; Alexa 568; red), and ATG14 (mouse anti-ATG14; Alexa 647; magenta) to examine the formation of the pro-bacterial autophagic initiation complex. GECs were then imaged via Leica DM6 CS Stellaris 5 Confocal/Multiphoton System at 63x. The Imaris software was used to obtain zoomed orthogonal views of (**Bi**) An infected GECs and (**Bii**) a theoretical autophagosome with HSp27, LC3C, ATG14, and Beclin 1 highly co-localized about it. The scale bar is 20 µm for all Magnification. All of the markers were found to have a Pearson correlation coefficient greater than .9 with each other via Imaris, denoting their close theorized interactions. Data is represented as Mean±SD, where n=3 and p<0.05 was considered as statistically significant via Student two-tailed T-test. *p<.05 **p<.005

### *P. gingivalis* induces specific pro-bacterial partnering of HSp27 and LC3C to promote its autophagic lifestyle

As LC3C-characterized, *P. gingivalis-*induced autophagy appears to be dependent upon HSp27 (**Fig. 3 and Fig. 4**) and Hsp27 and LC3C both appear to readily interact with one another to promote the maintenance of the Beclin 1-ATG14 pro-bacterial complex (**Fig. 8**), we wished to determine how HSp27 and LC3C may directly partner with one another during *P. gingivalis* infection. Confocal microscopy was used to show to specifically highlight how heightened LC3C readily co-localizes with increased levels of HSp27 during *P. gingivalis* infection in GECs, suggesting a direct pro-bacterial partnering (**Fig. 9A**; Pearson, r>0.88 for infected GECs; Two-Tailed Student T-Test, p<0.05). To confirm this theorized partnering, a far western approach was performed using full-length recombinant LC3C and HSp27 proteins (as we described previously for other molecules [41]). It was found that recombinant HSp27 actively binds to full-length LC3C in this assay format, suggesting that HSp27 partners with LC3C prior to LC3C’s lipidation as the pro-bacterial autophagosome forms (**Fig. 9B**). Importantly, further far western blots showed that HSp27 did not bind to either the LC3A or LC3B isoforms (**Fig. S4**). This lack of interactions confirmed the specificity of the partnering between HSp27 and the LC3C isoform during *P. gingivalis-*induced autophagy in infected GECs.

**Figure 9.**
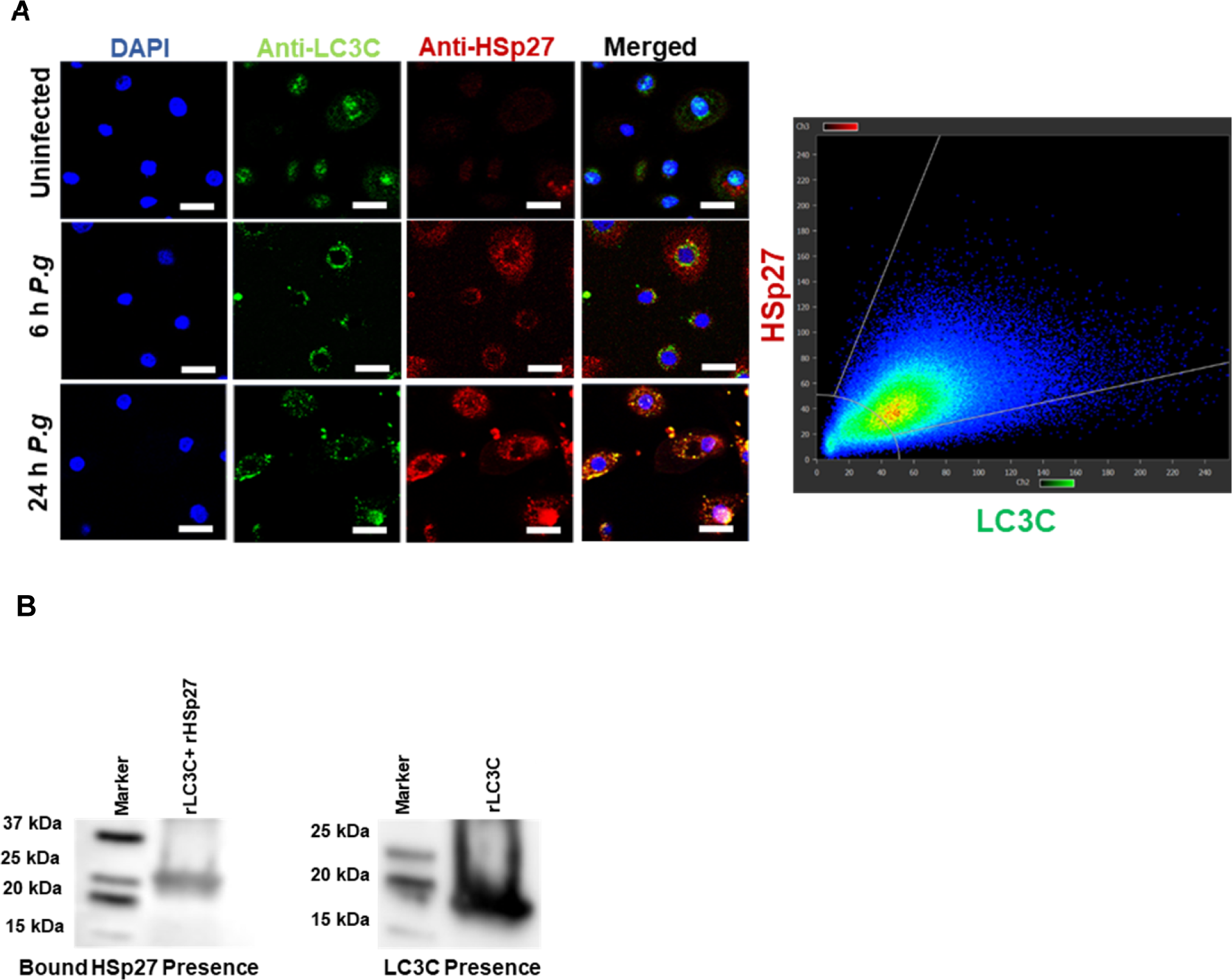
HSp27 and LC3C Selectively and Specifically Partner with One Another to Promote *P. gingivalis* (*P. g*)-Induced Autophagy. *P. g* was added at MOI 100 to Human Primary GECs, which were incubated 6 and 24 h. (**A**) GECs were stained for LC3C (rabbit anti-LC3C; Alexa 488; green) and HSp27 (mouse anti-HSp37; Alexa 568; red) following infection. HSp27 was found to readily and temporally colocalize with LC3C, having a Pearsons correlation coefficient of .85 at 24 h post infection via the Imaris post-processing software. (**B**) To assess if full length HSp27 is truly capable of binding to full-length LC3C, a far western approach was implemented by probing 5 µg of recombinant LC3C with 10 µg of recombinant HSp27 for one hour. Antibody specificity was accounted for via probing the LC3C blot with monoclonal mouse anti-HSp27 antibody (Not shown), which showed no cross-reactivity.

### Phosphorylated HSp27 (P-HSp27) preferentially binds to LC3C, inhibiting LC3C cleavage and halting the canonical maturation of *P. gingivalis-*specific autophagosomes

We then utilized various forms of *in silico* modelling to further elucidate the dynamics of the direct partnering between HSp27 and LC3C. Initially, Pymol was used to visualize the structures and functional regions of HSp27 and LC3C separately (**Fig. 10A and B).** Then, the two structural models were combined to identify specific theoretical polar and structural interactions between HSp27 and LC3C which verified our prior far western and confocal imaging findings indicating a direct physical interaction between these two molecules (**Fig. 10C**). Furthermore, given that HSp27 was previously established to be phosphorylated by the secreted Ndk effector of *P. gingivalis* on S78 and S82, the *in silico* models were made of LC3C combined with P-HSp27 to see how the additional phosphate group influenced the partnering (**Fig. 10D**). Interestingly, these models, which went through rigorous selection processes, strongly predicted that the engagement of LC3C with P-HSp27 resulted in the novel engagement of LC3C’s C-terminal tail, with two specific interactions forming between HSp27’s S78 and LC3C’s lysine (K)109 and HSp27’s S82 and LC3C’s arginine (R)106 (**Fig. 10D**). These interactions caused a shift in the predicted complex’s surface electrostatic potentials, induced greater buried surface areas, and resulted in higher interaction areas in the P-HSp27-LC3C complex (**Fig. 10E and F**). Interestingly, ATG4 did not appear in detectable levels in the mass spectrometry results of *P. gingivalis-*specific autophagosomes isolated from later points of infection (**Not Shown**), suggesting that the confirmational shift of LC3C caused by temporal P-HSp27-LC3C binding limited the interactions between ATG4 and LC3C.

**Figure 10.**
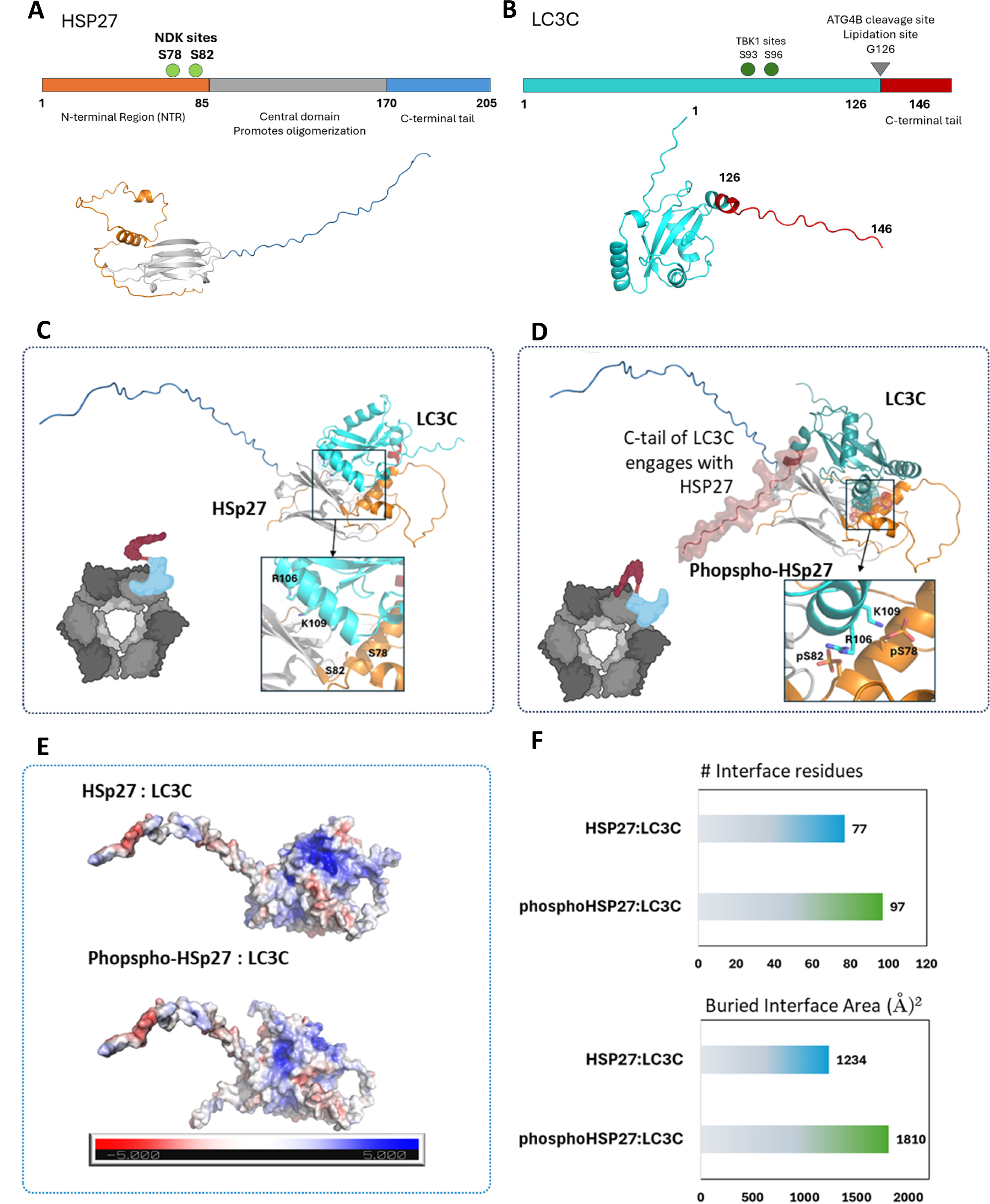
Phosphorylated HSp27 (P-HSp27) Preferentially Binds to the C-terminal Tail of LC3C, Inhibiting the Canonical Cleavage of LC3C and Halting the Canonical Maturation of LC3C-Specific Autophagosomes. The structural models of monomeric full-length wild-type (**A**) HSp27 (Uniprot: P04792) and (**B**) LC3C (Uniprot: Q9BXW4) were acquired from the AlphaFold database. Optimized complex configurations between (**C**) LC3C and unmodified HSp27 and (**D**) LC3C and P-HSp27 were then obtained, and the modifications in the theorized interaction sites in their N-terminal regions were highlighted. (**E**) Surface electrostatic potentials of the complexes were additionally mapped, contrasting the varied potentials between the two complexes. (**F**) Finally, the buried surface areas and interaction areas between HSp27 or P-HSp27 and LC3C proteins were also assessed and contact maps were generated.

Additional steps were then made to reconfirm the functional importance of Ndk on the *P. gingivalis* induced pro-bacterial autophagy machinery employing the isogenic *ndk* deletion mutant *P. gingivalis* strain (Δ*ndk P. gingivalis*) in biological assays with GECs. This mutant strain was previously shown to lack to induce a pro-survival phenotype in infected GECs [46] despite having the ability to internalize into GECs in comparable levels to the wild type strain of *P. gingivalis* [41,43]. Additionally, we discovered that Δ*ndk P. gingivalis* was intracellular survival deficient and underwent lysosomal trafficking as early as 6 hours post infection in GECs (**Fig. 11**). Thus, Δ*ndk P. gingivalis* was used to infect GECs alongside its wild-type counterpart to examine how the lack of Ndk-induced activation of HSp27 affected the complex formation between HSp27, Beclin 1, LC3C, and ATG14. It was identified that, unlike its wild-type counterpart, Δ*ndk P. gingivalis* did not induce the pro-bacterial complex assembly (**Fig. 12A**). Then, we performed phospho-mimicking HSp27 experiments with Δ*ndk P. gingivalis* infected GECs to see whether we could restore back the lost complex assembly and influence the intracellular fate of Δ*ndk P. gingivalis*. Transfection with the constitutively activated P-HSp27 construct was able to restore the HSp27-LC3C-Beclin 1-ATG14 pro-bacterial complex comparable to that seen through the usage of late-stage autophagic inhibitors (**Fig. 12A**) and was able to restore the live intracellular Δ*ndk P. gingivalis* levels (**Fig. S5**). Taken together, these results collectively support that the heightened phosphorylation of HSp27 is required for *P. gingivalis-*induced, pro-bacterial autophagy. Specifically, LC3C preferentially partners to P-HSp27, which can induce a confirmational shift to the C-terminal tail of LC3C, and potentially inhibit the final cleavage of lipidated LC3C’s tail by the ATG4 protease. This modification results in LC3C remaining tethered to the autophagosome, halting autolysosomal fusion (**Fig. 12B-C**).

**Figure 11.**
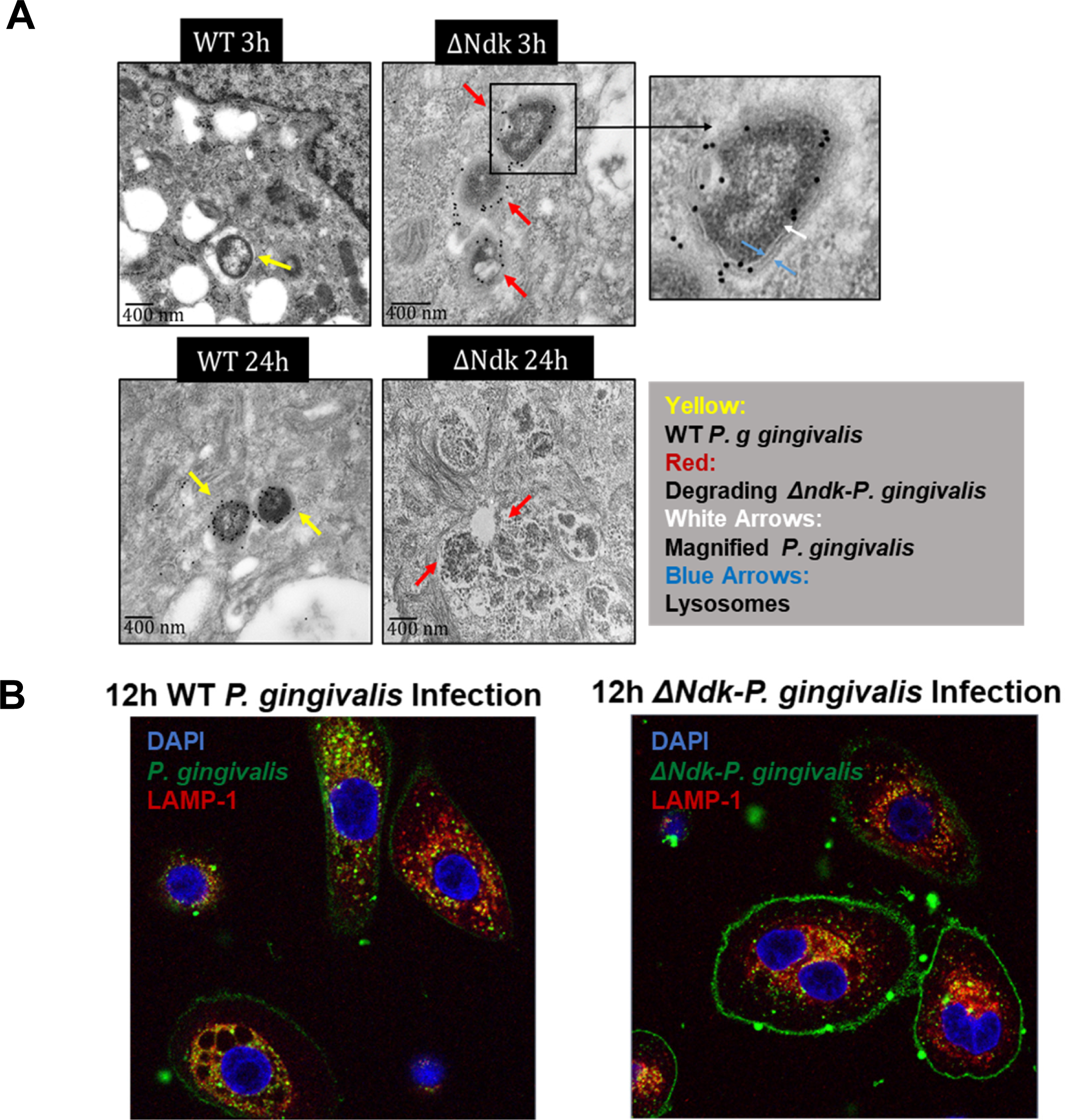
Δ*ndk-P. gingivalis (P. g)* Undergoes Canonical Degradative Autophagosomal Trafficking in Human Primary GECs. Primary GECs were infected with WT *P. gingivalis (P. g)* versus Δ*ndk*-*P. g* at MOI 100 for 3, 12, or 24h. (**A**) TEM analysis used Immunogold labeling for *P. g* was performed on WT *P. g* versus Δ*ndk*-*P. g* at 3 or 24h post-infection. Images were acquired at 40000x magnification utilizing a Hitachi H-7000 TEM (Hitachi High Technologies America, Inc.) affixed to a Veleta camera with iTEM. (**B**) Separately, GECs were additionally stained for *P. g* (rabbit anti-P*. gingivalis*; Alexa 488; green) and LAMP-1 (mouse anti-LAMP-1; Alexa 568; red) and were imaged. The range of all z-stacks was kept consistent and representative images were selected from the mid-ranged sections. Co-localization analysis of *P. g* and LAMP-1 was additionally carried out using the Zeiss LSM 880 Confocal Software, determining that the WT *P. g* had a Pearsons value of .137 while the Δ*ndk*-*P. g* had a Pearsons value of .704.

**Figure 12.**
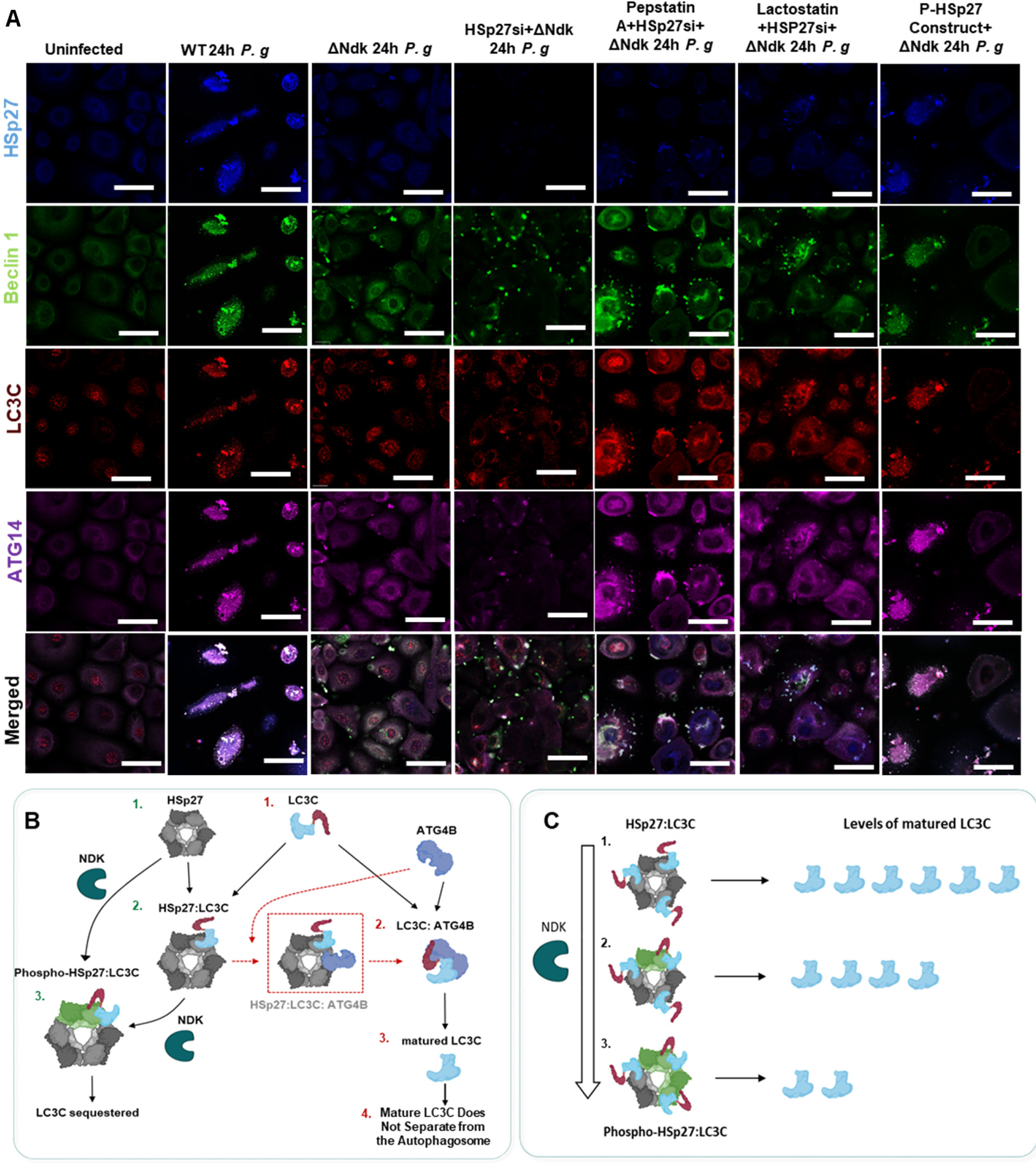
*P. gingivalis (P. g)* Secretes its Ndk Effector Molecule to Activate HSp27 and Induce Temporal HSp27-LC3C Partnering to Inhibit Canonical LC3C Cleavage by ATG4B and Halt Autolyosomal Fusion in GECs. A) Human primary GECs were treated with HSp27 siRNA (100nM) for 48 h or were transfected with 1 µg of the constitutively activated pFLAG-CMV2-HSP27-S78D/S82D construct for 48h. Select GECs were then jointly treated with the late stage autophagy inhibitors 1 µM Pepstatin A, or 1 µM lactostatin for 24h. Wild-type *P. g* or ΔNDK *P. g* was then added at MOI 100 to GECs, which were incubated for 24 h. (**A**) GECs also underwent staining for HSp27 (goat anti-HSp27; Alexa 405; blue), LC3C (rabbit anti-LC3C; Alexa 488; green), and Beclin 1 (sheep anti-Beclin 1; Alexa 568; red), and ATG14 (mouse anti-ATG14; Alexa 647; magenta) to examine the formation of the pro-bacterial autophagic initiation complex. GECs were then imaged via Leica DM6 CS Stellaris 5 Confocal/Multiphoton System at 63x. (**B**) A diagram was created detailing how LC3C preferentially partners to P-HSp27 over its non-phosphorylated counterpart, causing a confirmational shift to the C-terminal tail of LC3C. This shift results in the inhibition of the final lipidated LC3C tail cleavage by the ATG4B protease, lending to LC3C not disassociating from the autophagosome and halting fusion with the lysosome. (**C**) A diagram was also created to highlight the temporal relationship between HSp27 and LC3C, where LC3C can initially bind to HSp27 but via the actions of Ndk, it preferentially binds to P-HSp27, resulting in a limiting of mature, cleaved LC3C.

### The intracellular autophagic survival of *P. gingivalis* in a gingival-like environment significantly decreases after HSp27 depletion, in a manner mediated by oxidative stress

Provided these mechanistic findings with our reductionist human primary GECs, we wished to develop a further validation process where the cumulative effects of Hsp27 depletion could be highlighted in a more physiologically relevant, gingival-like environment. Thus, to replicate the epithelial junctions of the oral mucosa and examine the impact of HSp27 depletion on *P. gingivalis* colonization, a gingival organotypic culture system was constructed by Yilmaz lab with adaptations from previously described protocols [60,61] (**Fig. 13A**).

**Figure 13.**
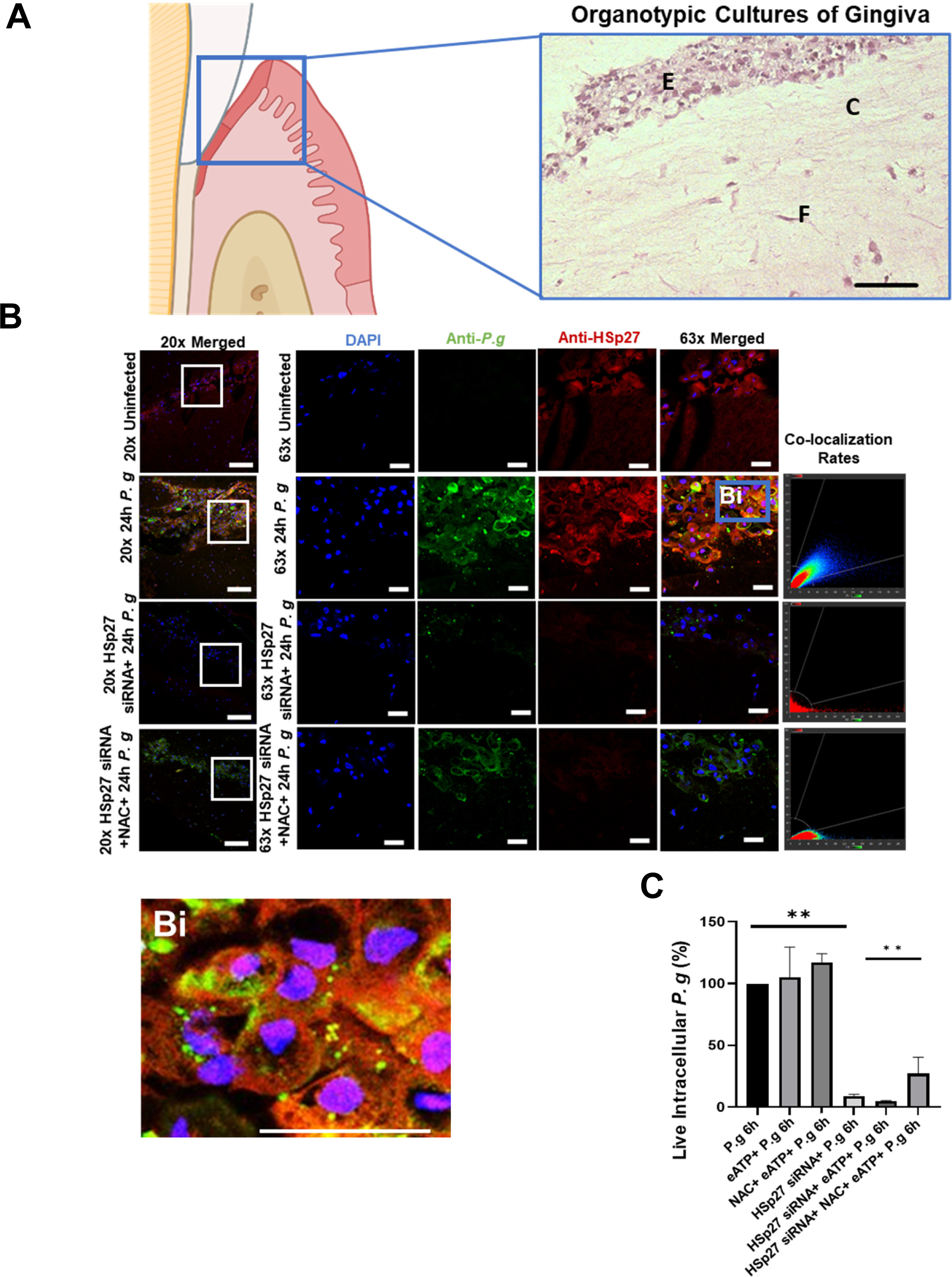
The Depletion of HSp27 Abrogates the Inhibition of Oxidative Stress and Severely Impacts the Intracellular Survival of *P. gingivalis (P. g)* Studied in Human Primary Organotypic Cultures of Gingiva. To create the organotypic culture systems, human primary GECs and Fibroblasts Cells (FBCs) were co-cultured together upon a collagen raft. Select rafts were then treated with HSP27 siRNA (100nM) for 48 h. Select rafts were also treated with N-acetyl Cysteine (NAC) (50 uM) for 1 h. *P. g* was added at MOI 100 to rafts, which were incubated for 24 h. Rafts were then collected and sectioned so that immunofluorescence could be performed. (**A**) Representative images of H&E stained raft culture systems at 20x magnification, which clearly mimic the oral gingival crevice. Scale bar is 50 µm. E: Multilayer Undifferentiated Epithelial Cells, C: Collagen Matrix; F: Fibroblasts. (**B**) Rafts were stained for *P. g* (rabbit anti-*P. g;* Alexa 488; green) or HpS27 (mouse anti-HSp27; Alexa 568; red). Rafts were then imaged via confocal microscopy (Leica DM6 CS Stellaris 5 Confocal/Multiphoton System) at 20x and 63x. Scale bar is 50 um for 20x and 20 um for 63x and Zoomed View. The range of z-stacks was kept consistent. HSp27 was once again found to readily co-localize with *P.g* with a Pearson coefficient of .83 as determined via the Imaris Software. (**Bi**) A Zoomed (4x) version of the 63x magnification was created using the Imaris Software, highlighting the intracellular nature of individual *P. g.* (**C**) GECs were additionally treated with HSp27 siRNA (100nM) for 48 h. Select GECs were then treated with N-acetyl Cysteine (NAC) (50 uM) for 1 h and/or eATP (3mM) for 30 min. *P. g* was added at MOI 100 to GECs, which were incubated 6 h. If any extracellular bacteria were present, they were killed by gentamicin (300 μg/mL) and metronidazole (200 μg/mL) treatment for 1 h. cDNAs were synthesized for qPCR using *P. g*-specific 16S rRNA primers to quantify intracellular level of live *P. g*. Data is represented as Mean±SD, where n=3 and p<0.05 was considered as statistically significant via One-Way Anova Test. ** p<0.005.

*P. gingivalis* was found to readily colocalize with induced HSp27 in the organotypic culture system (**Fig. 13B**), much akin to our prior human primary GEC monolayer findings. Upon depletion of HSp27 via targeted siRNA, there was marked impairment in *P. gingivalis* colonization levels of the primary GEC multi-layer and the underlying embedded human primary fibroblast cells (FBCs) (**Fig. 13B**). Treatment with NAC, however, was able to partially restore *P. gingivalis* levels in HSp27 depleted organotypic system (**Fig. 13B**). These findings were then confirmed through the usage of an *in situ*, *P. gingivalis*-specific, antibiotic protection assay. Our analyses found that intracellular *P. gingivalis* levels were significantly diminished in HSp27-depleted GECs, cementing HSp27’s relevance in prolonged *P. gingivalis* survival in the oral mucosa (**Fig. 13C**). The treatment of HSp27-depleted, infected cells with eATP further decreased the intracellular bacteria levels, which were partially restored via treatment with NAC, once again underlying the key role of HSp27 in cell redox homeostasis to facilitate the autophagic survival of *P. gingivalis* in GECs.

### Cross-sectional human ex-vivo analyses support high expression levels and increased co-localization of *P. gingivalis*, HSp27, and LC3C in chronically inflamed oral tissues

Given the findings of the organotypic culture systems, excised human gingival tissues were also used to study HSp27 and LC3C relevance in *P. gingivalis*-induced diseases. Initially, we did so via analyzing the results from a bioinformatics study published on the NCBI (Gene Expression Omnibus) GEO database. From that study, LC3C and HSp27 mRNA expression levels were measured in periodontitis-afflicted and healthy periodontal tissues. We found that both HSp27 and LC3C expression levels were induced in periodontal-diseased tissues, when compared to their healthy counterparts (**Fig. 14A and 14B**). Additionally, we performed immunofluorescence staining on human gingival epithelium specimen. In doing so, we discovered that there were substantially higher (∼4.5 fold) levels of *P. gingivalis* in periodontitis-afflicted tissue when compared to healthy counterparts (**Fig. 14C and Fig. S6A**); which is consistent with prior publications [10]. Additionally, however, both LC3C and HSp27 levels were also found to be significantly increased (∼5 fold and ∼2 fold respectively) in periodontitis-afflicted tissues (**Fig. 14D and Fig. S6B and S6C**). LC3C and HSp27 exhibited higher levels of co-localization with each other and *P. gingivalis* (Pearson coefficient of 0.85 for LC3C and *P. gingivalis* and 0.93 for LC3C and HSp27) (**Fig. 14C and Fig. 14D)** than their healthy counterparts. These cross-sectional findings provide visual and statistical information indicating the relevance of HSp27 and LC3C in *P. gingivalis-*driven diseases, and once more pinpoint the close partnering of these molecules induced by heightened *P. gingivalis* presence.

**Figure 14.**
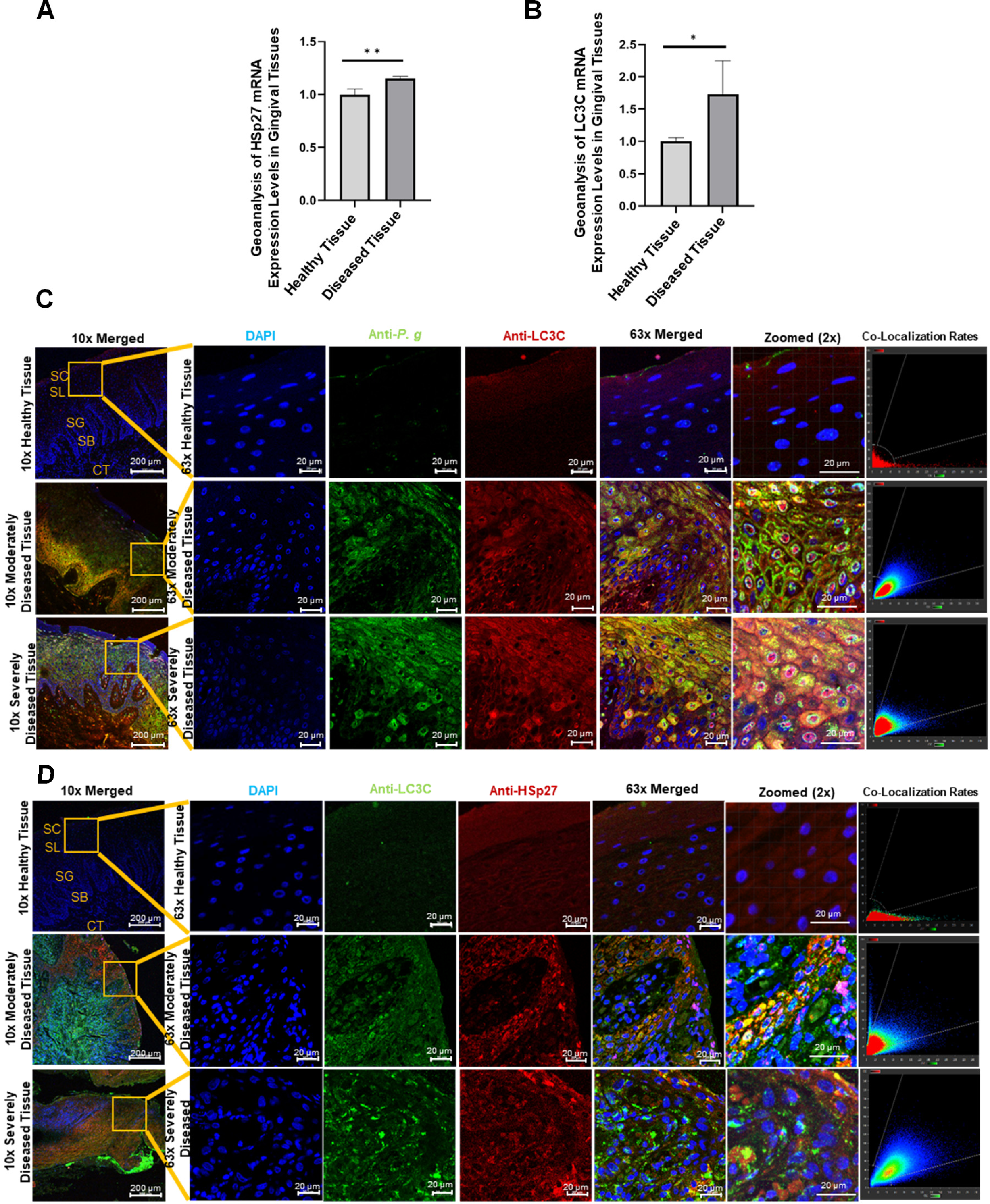
Cross-Sectional Human *in Situ* Sample and Expression Analyses Support High Levels and Increased Co-localization of *P. gingivalis (P. g)*, HSp27, and LC3C in Periodontitis-Afflicted Oral Tissues. Publicly available mRNA expression data (GEO accession: GSE79705) was obtained from previously collected and examined periodontitis-afflicted and healthy gingival tissues. This microarray expression data was then analyzed via GEO2R and the relative levels of (**A**) HSp27 and (**B**) LC3C were obtained and compared. Data is represented as Mean±SD, where n=12 and p<0.05 was considered as statistically significant via One-Way Anova. *p<0.05. Representative confocal images of gingival biopsy specimens from healthy individuals and periodontitis-afflicted patients were also taken and examined. DAPI staining was utilized to visualize cellular DNA. (**C**) *P. g* (mouse anti-P*. gingivalis*; Alexa 488; green) and LC3C (rabbit anti-LC3C; Alexa 594; red) were detected via dual staining. (**D**) HSp27 (mouse anti-HSp27; Alexa 488; green) and LC3C detection (rabbit anti-LC3C; Alexa 594; red) were also detected. Images were then captured using super resolution confocal laser scanning microscopy (Leica DM6 CS Stellaris 5 Confocal/Multiphoton System) at 10x and 63x magnification with oil immersion. Zoomed (2x) 3D versions of the 63x magnifications were obtained via the Imaris software. The range of z-stacks was kept consistent. SC: Stratum corneum, SL: Stratum lucidum, SG: Stratum granulosum, SB: Stratum basale, LT: Lamina propria. Scale bars = 200µm for 10x and 20 µm for 63x. Quantification of mean fluorescence intensity provided in Supplement. LC3C and HSp27 both were found to exhibit high levels of co-localization with each other and with *P. g,* as the Pearson coefficient was calculated to be .93 for LC3C and *P. g* and .85 for LC3C and HSp27 via the Imaris Software at the most severe state of disease.

## DISCUSSION

Autophagy is an essential process that maintains homeostasis and over-all cellular health. During selective autophagy, following its activation, the Beclin 1 Core Complex joins with ATG14 to induce the formation of ATG14-Involved Initiation Complex [20,24,34]. This complex then induces autophagosomal nucleation and the sequestration of damaged cellular components and foreign bodies within double-membraned autophagosomes [20,24,34]. These autophagosomes are then trafficked to lysosomes, where they undergo autolysosomal fusion and resulting degradation [24,34]. A variant of autophagy, dubbed xenophagy, specifically targets and removes invading bacteria from host cells [35]. However, like certain other noted bacterial species, *P. gingivalis* appears to be able to evade autophagic recognition and even usurp the autophagic protective host mechanisms in order to enhance its own survival in a variety of cell types, including GECs [15,62]. However, the autophagic machineries that facilitate this have largely been remained elusive. The novel findings of this paper expound upon these processes, detailing how, following *P. gingivalis’* invasion of GECs, the bacteria induce HSp27 and LC3C to promote a unique non-canonical, pro-bacterial form of autophagy. HSp27 is activated and induced by *P. gingivalis-*infection, following which, HSp27 closely regulates *P. gingivalis-* driven pro-bacterial autophagy via the upstream partnering with the key selective autophagy molecule, LC3C, prior to its incorporation during phagophore nucleation and autophagosomal formation. Upon gradually increasing phosphorylation of HSp27 on S78 and S82 by the bacterial Ndk [42], P-HSp27 preferentially and tightly partners with LC3C, modifying the placement of the LC3C C-terminal tail, which could hinder the ATG4 protease from further cleaving LC3C following LC3C conjugation with PE and incorporation into the autophagosomal membrane. This inaccessibility appears to render LC3C remain incorporated into the autophagosomal membrane. Thus, LC3C and P-HSp27 stay partnered and accumulate on the autophagosomal membrane. As LC3C cannot disassociate, *P. gingivalis-*specific autophagosomes cannot fully mature and do not fuse with lysosomes. This continual immaturity is exemplified by the apparent sustained binding of the autophagic initiation molecules Beclin 1 and ATG14, where the canonically transient ATG14-Involved Initiation Complex appears to become firmly tethered to *P. gingivalis*-specific autophagosomes. It is tempting to speculate that the joint partnering of LC3C and P-HSp27 with these two autophagic initiation molecules alters the formation of the ATG14-Involved Initiation Complex, causing the complex to become constantly locked onto the autophagosomes. This structural shift could also make it so that other autophagic maturation modulators, such as UV irradiation resistance-associated gene (UVRAG), are not granted the chance to competitively bind to the Beclin 1 Core Complex [26, 27]. This theory is strengthened by the mass-spectometry and western blot analyses of isolated *P. gingivalis-*specific autophagosomes, where UVRAG was not detectable (**Not Shown**). Such results provide further specificity for how *P. gingivalis-*specific autophagosomes do not canonically mature, undergo autolysosomal fusion, or undergo lysosomal acidification.

Concurrently, the non-canonical, pro-bacterial lifestyle of *P. gingivalis* appears to be influenced and interlinked with other infection-modulated cellular pathways, such eATP generation and the resulting oxidative stress. One of the main host aspects that *P. gingivalis* must overcome to prolong its intracellular variant of survival is the potent pro-inflammatory release of small danger molecules, such as eATP, caused by the GEC’s response to bacterial infection [43,56,63,64]. It has been well-characterized that, following bacterial invasion of GECs, eATP is generated and released by infected cells, promoting H_2_O_2_ formation [42,45,65]. *P. gingivalis,* however, has been shown to mitigate antibacterial NADPH oxidase-mediated H_2_O_2_ generation and HOCl production via influencing the host-antioxidant GSH synthesis [42]. Interestingly, prior studies outside of host-microbe field have also shown that HSp27 is capable of inducing GSH synthesis [47,48,66]. Thus, this study aimed to highlight this potential pro-bacterial HSp27-GSH interaction and attempted to define its importance in *P. gingivalis-*induced non-canonical autophagy. During the prolonged intracellular survival of *P. gingivalis* in GECs, the *P. gingivalis-*specific autophagosomes were found to be readily enriched in GSH. Additionally, GSH levels were induced in a manner dependent upon HSp27 presence. Finally, the inductions of LC3C, and ATG14 / Beclin 1—the major emerging hallmarks of *P. gingivalis-*induced, non-canonical autophagy were increasingly disrupted following eATP treatments of HSp27-depleted GECs. For these reasons, it is justified to theorize that HSp27 is jointly inducing GSH synthesis to combat the anti-bacterial redox stress and maintain a pro-bacterial redox environment within the infected GECs. The effects seen during HSp27 depletion lend credence to this theory: if HSp27 was absent in infected GECs, the integrity of the *P. gingivalis*-induced, non-canonical autophagy collapsed. It is only when NAC, a major GSH precursor, was added to the HSp27-depleted GECs, that the pro-bacterial autophagic flux and interactions were capable of being even partially rescued. It is plausible that novel GSH was thus being synthesized independent of HSp27 presence, restoring a partially favorable cellular environment for *P. gingivalis* survival. Without GSH-synthesizing alternatives, the unabrogated oxidative stress overtakes the infected GECs and anti-bacterial canonical autophagy is induced once more, killing intracellular *P. gingivalis*.

Our novel study highlights, for the first time, how activated HSp27 specifically partners with LC3C to inhibit its canonical maturation, which impacts the formation of the Beclin 1-ATG14-specific nucleation complex, resulting in a unique form of *P. gingivalis-*induced, non-canonical, pro-bacterial variant of autophagy (**Fig. 15**). These interactions are closely interconnected to the redox-state of the infected host cells, as HSp27 appears to be separately inducing GSH synthesis to mitigate the impact of eATP-induced ROS production. These new findings underpin how HSp27 and its partnering with LC3C could be vital autophagic molecular targets involved in the intracellular survival of *P. gingivalis* in GECs. Identifying the interactors involved in the usurpation of host autophagic mechanisms is critically important, as the intracellular lifestyle of *P. gingivalis* is recalcitrant to conventional antimicrobial control procedures and *P. gingivalis* appears to establish repeated long-term infections in infected GECs [67]. The *in silico* models identified a number of strong theoretical interaction sites between P-HSp27 and LC3C. Of these sites, seven appeared to be highly specific to the LC3C isomer— given that LC3A and LC3B were not seen to exhibit direct partnering with HSp27 via far western analyses (**Table 1 and Fig. S4**). Future studies will focus upon these putative interaction sites to see if mutating them hampers LC3C and P-HSP27 binding and the autophagic life of *P. gingivalis.* Such therapeutic targeting could be highly effective, as these interactions appear to be very specific for the autophagic lifestyle of *P. gingivalis* in GECs. Future work will also be performed to further elucidate how HSp27 and LC3C partnering could also impact other theorized autophagic interactions, such as the inactivation of UVRAG and other maturation modulators.

**Figure 15.**
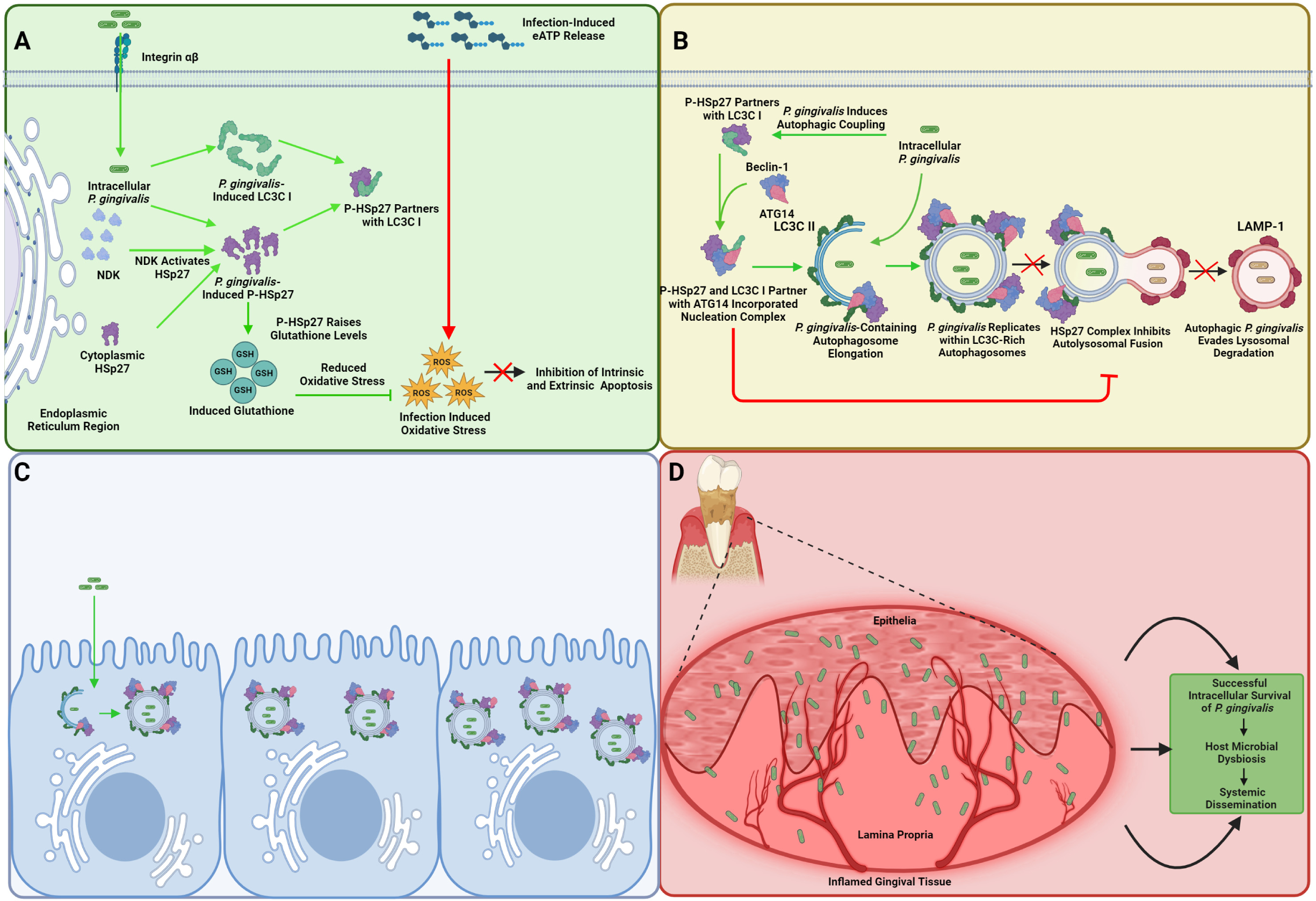
HSp27 is a Critical Regulator in Pro-bacterial LC3C-Characterized Autophagy, Facilitating the Intracellular Autophagic Survival of *P. gingivalis (P. g)* and Influencing the Bacterial Symbiosis of the Oral Mucosa. The proposed diagram of the identified mechanisms of *P. gingivalis* persistence in GECs. (**A**) HSp27 is largely induced and spatially recruited by *P. g* invasion of the host cells. After the initial periods of cellular infection, the gradually growing secretion of the bacterial Nucleoside-diphosphate-kinase (Ndk) into the cytoplasmic space causes heightened activation of HSp27 (P-HSp27) via direct phosphorylation. P-HSp27 abrogates extracellular ATP (eATP)-induced antimicrobial Reactive-Oxygen-Species (ROS) production via increasing glutathione (GSH) levels. In parallel, *P. g*-mediated induction of HSp27 promotes the specific recruitment and lipidation of LC3C, an isomer of the LC3 autophagosomal structural molecule, which is strictly dependent upon the large presence and the strong antioxidant activity of HSp27. LC3C and HSp27 partner in a stepwise manner, 1) their coupling drives the formation of Beclin1/ATG14 induction, 2) where the temporally increased phosphorylation of HSp27 by *P. g* Ndk both strengthens the P-HSP27 and LC3C partnering and shifts the confirmation of the LC3C tail so that LC3C cannot be successfully further cleaved by ATG4. (**B**) P-HSp27 and LC3C become increasingly assembled to the ATG14-Incorperated Nucleation Complex, where they prolong the complex’s formation and result in the accumulation of the complex on forming autophagic membranes. Thus, the strengthened partnering between P-HSp27 and LC3C is the proposed mechanism for inhibiting the autolysosomal fusion of *P. g-*specific autophagosomes, which is also controlled by the host cell redox homeostasis (**C**) Autophagic *P. g* does not undergo lysosomal degradation and is instead able to survive, multiply and subsequently intercellularly spread to neighboring cells to propagate. (**D**) Thus, the non-canonical, pro-bacterial autophagic events create a favorable and protected cellular environment for *P. g*, thereby establishing long-term intracellular bacterial persistence. The chronic colonization of *P. g* in the epithelia can lead to host-microbial dysbiosis in oral mucosa and systemic disorders.

**Table 1.** *P. gingivalis (P. g)* Induces HSp27 and LC3C Partnering to Promote its Autophagic Survival in GECs. (**A**) Interfacing residues between HSP27 and LC3C in the HSP27:LC3C modelled complex. (**B**) Major putative interaction sites of the HSp27-LC3C complex were compared to LC3A and LC3B, denoting that the following putative interactions are specific only to HSp27 and LC3C: 1) the Gln5 in the N-terminus of LC3C interacts closely with both the Trp42 and Glu41 of HSp27, 2) LC3C’s Arg46 interacts with both Ser50 and Trp45 of HSp27, 3) LC3C’s Arg80 interacts with Gly34 of HSp27, 4) LC3C’s Asn91 and Ala132 exhibit significant interactions to HSp27’s Tyr54, 5) LC3C’s Glu129 interacts with Arg56 of HSp27, 6) LC3C’s Pro133 interacts with HSp27’s Leu58, and finally, 7) Arg142 of LC3C exhibits significant polar bonding with Glu64 of HSp27.

| HSP27 | LC3C |
| --- | --- |
| A:SER 15 | C:PRO 4 |
| A:TRP 16 | C:GLN 5 |
| A:PRO 18 | C:LYS 6 |
| A:PHE 19 | C:ILE 7 |
| A:ASP 21 | C:SER 9 |
| A:ARG 37 | C:PRO 12 |
| A:LEU 38 | C:LYS 14 |
| A:GLU 40 | C:GLN 15 |
| A:SER 43 | C:SER 18 |
| A:SER 65 | C:LEU 19 |
| A:PRO 66 | C:ALA 20 |
| A:ALA 67 | C:ILE 21 |
| A:VAL 68 | C:GLN 23 |
| A:ALA 70 | C:VAL 26 |
| A:PRO 71 | C:ALA 27 |
| A:ALA 72 | C:ARG 30 |
| A:SER 74 | C:ALA 31 |
| A:ARG 75 | C:PRO 34 |
| A:SER 78 | C:GLU 42 |
| A:ARG 79 | C:TYR 44 |
| A:SER 82 | C:PRO 45 |
| A:VAL 85 | C:ARG 46 |
| A:GLU 87 | C:ASN 90 |
| A:ARG 89 | C:ASN 91 |
| A:ASP 100 | C:LYS 92 |
| A:SER 156 | C:TYR 105 |
| A:SER 158 | C:ARG 106 |
| A:PRO 159 | C:ASP 107 |
| A:GLU 160 | C:TYR 108 |
| A:THR 162 | C:LYS 109 |
| A:THR 164 | C:ASP 110 |
|  | C:GLU 111 |
|  | C:ASP 112 |
|  | C:GLY 113 |
|  | C:TYR 116 |
|  | C:THR 118 |
|  | C:LEU 128 |
|  | C:GLU 129 |
|  | C:ALA 131 |
|  | C:ALA 132 |
|  | C:PRO 133 |
|  | C:ASP 135 |

**B** **Major Interacting amino acid residues between HSP27 and LC3C**
| Amino acids in HSP27<sup>a</sup> | Amino acids in LC3C<sup>b</sup> | Amino acids in LC3A/LC3B* |  |
| --- | --- | --- | --- |
|  |  | LC3A<sup>c</sup> | LC3B<sup>d</sup> |
| F (Phe) 33 | T (Thr) 82 | T (Thr)76 | N (Asn)76 |
| G (Gly) 34 | R (Arg) 80 | N (Asn) 74 | N (Asn) 74 |
| E (Glu) 41 | Q (Gln) 5 | Absent | Absent |
| W (Trp) 42 | Q (Gln) 5 | Absent | Absent |
| W (Trp) 45 | R (Arg) 46 | G (Gly) 40 | G (Gly) 40 |
| S (Ser) 50 | R (Arg) 46 | G (Gly) 40 | G (Gly) 40 |
| Y (Tyr) 54 | A (Ala) 132 | Absent | Absent |
| Y (Tyr) 54 | N (Asn) 91 | Q (Gln) 85 | G (Gly) 85 |
| Y (Tyr) 54 | S (Ser) 93 | S (Ser) 87 | S (Ser) 87 |
| R (Arg) 56 | E (Glu) 129 | Absent | L (Leu) 123 |
| L (Leu) 58 | P (Pro) 133 | Absent | Absent |
| E (Glu) 64 | R (Arg) 142 | Absent | Absent |

\*Most of the amino acids are either different or missing in LC3A/LC3B compared to those in LC3C and that explains Far western result why LC3A/LC3B don't interact with HSP27.
Protein IDs used for analysis: <sup>a</sup>HSP27: P04792, <sup>b</sup>LC3C: Q9BXW4, <sup>c</sup>LC3A: Q9H492, and <sup>d</sup>LC3B: Q9GZQ8

*P. gingivalis* has recently been postulated to participate in multiple chronic systemic diseases etiologies and progression, such as Alzheimer Disease, oro-digestive cancers, and Rheumatoid Arthritis [2,4,5,68]. Because of these associations, studies targeting activated HSp27 coupled with pro-bacterial autophagic mechanisms could not only result in effective therapeutic strategies that control *P. gingivalis’* chronic colonization of the oral cavity, but also could potentially curtail the pathologies of associated with these devastating co-morbid chronic diseases. Additionally, further studies examining the non-canonical autophagic mechanisms *P. gingivalis* influences to survive could abet in curtailing or targeting other pathogenic bacteria that usurp host autophagic mechanisms to prolong their own intracellular survival.

## MATERIAL AND METHODS

### Bacteria Culture

*P. gingivalis* ATCC 33277 and its isogenic Δ*ndk* mutant were anaerobically cultured in trypticase soy broth (TSB) supplemented with yeast extract (1 mg ml^−1^), hemin (5 µg ml^−1^) and menadione (1 µg ml^−1^) at 37°C [12,14,17,43]. Erythromycin (10 mg/ml) was added to the TSB media as a selective agent for the mutant strain’s growth as previously described [45]. *P. gingivalis* was cultured overnight and was then centrifuged at 6000 g for 10 minutes at 4 °C. Following harvest, *P. gingivalis* resuspended in Dulbecco’s phosphate-buffered saline (PBS) (HyClone). Following resuspension, the total number of bacteria at mid-long phase were calculated utilizing a Klett-Summerson photometer. It has been shown that the usage of an inoculum of MOI 100 is optimal for the attachment and invasion rate of *P. gingivalis* in primary GECs; thus, MOI 100 was utilized throughout this study [12,14,17].

### Primary Cell Culture

Human primary gingival epithelial cells (GECs) were obtained following the oral surgery of adult patients who were accepted for either tooth crown lengthening procedures or impacted third molar extraction. Following the randomization and anonymization of the patients, under the approved guidance of the University of Florida Health Science Center Institutional Review Board (IRB, human subjects assurance number FWA 00005790), healthy gingival tissue samples were collected as previously described [55]. Following isolation, GECs were cultured in serum-free keratinocyte growth medium (KGM, Lonza, 10130685) at 37°C in a humidified 5% CO_2_ incubator. Starvation conditions were induced via incubation in HBSS (ThermoFisher Scientific, 14025092) for 6 h prior to collection.

Human primary fibroblast cells (FBC) were obtained and isolated via the same procedures listed for GECs. Following isolation, FBCs were grown in Fibroblast Basal Medium (ATCC, PCS-201-030) with ATCC® Fibroblast Growth Kit-Low Serum (ATCC, PCS-201-041) supplements added.

### Organotypic Raft Culture

Once primary GECs and FBCs initially reached 80% confluence, the FBCs were then trypsinized and centrifuged at 59 x g for 5 min. Simultaneously, a collagen raft mixture was prepared on ice via combining 3 mg/ml collagen with 10X E-Media (Powdered DMEM (Millipore Sigma, D7777-10L), powdered Ham’s F-12 (Millipore Sigma, N6760), and di H_2_O) and Reconstitution Buffer (2.2% NaHCO3, .15N NaOH, 200 mM HEPES (Millipore Sigma, H3375), and di H_2_O). Tyrpsin was then decanted, FBS was added to the FBCs at 0.2 mL/raft, the FBCs were added to the collagen raft mixture, and the FBC raft mixture was added to the bottom of 35 mm dishes. The plates were then placed at 37°C in 5% CO_2_ and were allowed to congeal. Primary GECs were then seeded atop the collagen in KBM media. The rafts were refed daily with KBM for 6 days, and were disturbed daily via tapping to prevent attachment. The rafts were then raised via being transferred onto a 50 mm raised wire disk (McMaster Carr, 4195A11) and placed into a 100 mm dish. After raising, the rafts were refed daily and were permitted to grow for another 4 to 6 days, pending visible growth. Following HSp27 siRNA transfection and *P. gingivalis* infection, rafts were washed with PBS and were placed in 5 mL of 10% Neutral buffered formalin (NBF) for 18 h prior to being transferred to 70% EtOH until paraffinization.

### SiRNA Knockdown

For HSp27 depletion experiments and LC3C deletion experiments, GECs were treated with HSp27-specific siRNA (100 nM, Cell Signaling Technology, 6356S) or LC3C-specific siRNA (100 nM, ThermoFisher Scientific, 4392421) in RNAiMax Lipofectamine transfection reagent (Thermofisher Scientific, 13778150) for 48h as per company protocol.

### Oxidative Stress Treatments

For HSp27 depletion experiments and LC3C deletion experiments, GECs were additionally treated with N-acetyl Cysteine (NAC) (50 uM, Millipore Sigma, A9165) for 1 h and/or eATP (3mM, Millipore Sigma, A6419) for 30 min prior to 6, 12, or 24 h infection with *P. gingivalis*. For proteomics analyses, GECs were treated with buthionine sulfoximine (BSO) (100 nM, Millipore Sigma, B2515-500MG) for 48h prior to 3, 6, or 24h infection with *P. gingivalis.* Additionally, the viability of the cells following these specific treatments was tested via MTT assay, to confirm the lack of cellular toxicity in the optimized concentrations (not shown).

### Autophagic Inhibitor Treatments

Select cells were treated with pepstatin A (1 µM, Millipore Sigma, P2032), 3-Methyladenine (5 mM, Millipore Sigma, M9281), lactostatin (1 µM, Millipore Sigma, L5670) or Bafilomycin A1 (1 µM, Millipore Sigma, 19-148) for 24 h prior to siRNA knockdown, oxidative stress treatments, and infection with *P. gingivalis* at an MOI of 100 [52–54].

### mCherry-eGFP-LC3C Construct Construction

This mCherry-eGFP-LC3C construct has an LC3C conjugated to an acid insensitive mCherry (red) probe and an acid sensitive eGFP (green) probe, which allow for the visualization of autophagolysosomal fusion and the resulting acidification as the eGFP probe is quenched. To create this construct, at first the coding sequence of LC3C (444 nt) from mRNA sequence (GenBank Accession # NM_001004343.3) with addition of EcoR1 and SalI restriction sites was synthesized by IDT DNA Technologies (Coralville, IA). Next, the LC3B in pBABE-puro mCherry-EGFP-LC3B (Addgene, Plasmid #22418) was replaced with the LC3C at EcoRI and SalI site to make pBABE-puro mCherry-EGFP-LC3C. At the same time, an empty vector without LC3C was constructed by removing LC3B from pBABE-puro mCherry-EGFP-LC3B and religating the pBABE-puro mCherry-EGFP part. The pBABE-puro mCherry-EGFP-LC3C and empty vector pBABE-puro mCherry-EGFP were confirmed by restriction digestions with EcoR1-SalI and NaeI-HindIII, respectively, as well as by sequencing (Gene Universal, Newark DE).

### mCherry-eGFP-LC3C Transfection

GECs were transfected with 1 µg of mCherry-eGFP-LC3C construct plasmid or a control plasmid (empty vector without LC3C) for 48 h using Lipofectamine 3000 (Thermofisher Scientific, L3000001) in serum free KGM medium as per the manufacturer’s instructions.

### pFLAG-CMV2-HSp27-S78D/S82D Transfection for Phospho-HSp27 Mimicking

GECs were transfected with 1 µg of pFLAG-CMV2-Hsp27-S78D/S82D (Addgene plasmid # 85187 [69]), a constitutively activated HSp27 construct, for 48 h using Lipofectamine 3000 (Thermofisher Scientific, L3000001) in serum free KGM medium as per the manufacturer’s instructions. This plasmid was provided as a gift from Ugo Moens.

### 16S RNA-based Antibiotic Protection Assays

The intracellular invasion and survival of *P. gingivalis* in primary GECs was determined as we previously described in. In brief, GECs were plated at a density of 2.5 × 10^5^ in 6 well plates and were cultured until ∼80% confluence. Following select wells undergoing siRNA transfection or eATP treatments, and *P. gingivalis* infection, GECs were washed in PBS and were incubated in gentamicin (300 µg/mL) and metronidazole (200 µg/mL) for 1 h to remove any extracellular bacteria [55]. Following antibiotic treatment, the total RNA of each condition was isolated and collected using Trizol Reagent (Invitrogen, 15596026). Any genomic DNA contamination was removed via DNase treatment (Ambion, AM2222). Following DNase digestion, cDNA was synthesized utilizing 1 μg of each condition’s total isolated RNA via a High-Capacity cDNA Reverse Transcriptase Kit (Applied Biosystems, 4368814). A 1:10 dilution of cDNA was used to detect *P. gingivalis*-specific 16s rRNA via SYBR Green Real-time qPCR (Forward: 5 -TGTAGATGACTGATGGTGAAAACC-3 ; Reverse:5 -ACGTCATCCCCA CCTTCCTC-3). qPCR was then conducted via the CFX96 real-time system (Bio-Rad, 1845097), with a cycle of 98°C for 3 min followed by 40 cycles of 9 5°C for 15 s, 60.7°C for 30 s, and 72°C for 30 s. A standard curve was created, as described in [10,55]. This standard curve was used to determine the colony forming units (CFU) of corresponding Ct (Threshold cycle) values obtained via qPCR analysis.

### The Magnetic Isolation of *P. gingivalis-*Specific Autophagosomes

*P. gingivalis* ATCC 33277 strain was incubated in 5 μg/mL Lipobiotin (LB) in PBS overnight at 4°C. Labelled *P. gingivalis* was then incubated with MagCellect Streptavidin Ferrofluid (R&D Systems, MAG999) at a concentration of 7–14 × 10^8^ particles per 2 × 10^8^ bacteria for 20 min at 4°C. To determine the importance of HSp27 on *P. gingivalis*-specific autophagosomal integrity, GECs were plated at a density of 1.4 × 10^6^ in 100 mM dishes and were cultured until they reached ∼80% confluence. Following siRNA transfection, redox-related treatments, and *P. gingivalis* infection, *P. gingivalis*-specific autophagosomes were then isolated as described in [50]. In brief, GECs were washed with Buffer A (15 mm HEPES buffer with 20 mm sucrose, 50 mm MgCl_2_, and protease inhibitors containing 0.02% ethylenediaminetetraacetic acid (EDTA)) to remove non-ingested bacteria, were trypsinized, and resuspended in Buffer A combined with Benzonase Nuclease (25 U/mL, Millipore Sigma, E1012-25KU) and Cytochalasin D (5 µg/mL, Millipore Sigma, C8273). Following resuspension, cell membranes were disrupted and *P. gingivalis*-specific autophagosomes were released via sonicating GECs on ice via a ThermoFisher Qsonica Sonicator Q500 (Fisher Scientific, 15-338-282) at an amplitude of 75% for 10 s. GECs were then centrifuged at 300 g for 2 min between sonication steps, after which the supernatant was removed and replaced with 500 μL of Buffer A. After the third sonication, the samples were left on ice for 10 min. The supernatants were then removed and incubated at 37°C for 5 min. Following incubation, the samples were placed in a 16-Tube SureBeads™ Magnetic Rack (Bio-Rad, 1614916) for 30 min prior to repeated manual rinsing of the magnetic fraction with Buffer A at a flow rate of 250 μL/min for 15 min to ensure purity. Finally, the autophagosome-containing fractions were collected and centrifuged at 15,000 g for 12 min, before the supernatant was decanted and the autophagosomes were resuspended in 100 μL of HEPES buffer combined with protease inhibitors.

### Proteomic analysis of *P. gingivalis-*Specific Autophagosomes

The enriched proteins of isolated *P. gingivalis-*specific autophagosomes were analyzed via label-free quantification (LFQ) by LC-MS/MS using Orbitrap Fusion Lumos with ETD/UVPD, a high resolution and high mass accuracy instrument (Thermo Fisher Scientific) that is maintained and performed by the Proteomics Core at MUSC. The levels of HSp27 (Protein ID# P04792), LC3 (Protein ID# H3BTL1), Beclin 1(Protein ID# O14964), and ATG14 (Protein ID# Q6ZNE5) were quantified based on the peak intensity of the corresponding MS spectrum of HSp27, LC3, Beclin 1, and ATG14 with 95% confidence. The specificities of the protein identifications were confirmed via BlastP searches using individual peptide fragments of each of the proteins. At least six replicates were pooled together in each assay.

### Western Blotting Analyses

Following siRNA transfection, eATP and NAC treatments, and *P. gingivalis* infection, GECs either underwent the selective isolation of *P. gingivalis-*specific autophagosomes or were lysed with .2% Triton-X lysis buffer combined with protease and phosphatase inhibitors: 1mM PMSF; 1mM NaF; Halt Phosphatase inhibitor cocktail (ThermoFisher, 78430); and Protease Inhibitor Cocktail Set III (Millipore Sigma; 535140). Equal amounts of the samples’ total protein were determined by Bradford Assay, samples were subjected to 4x Laemmli sample buffer (BioRad, 1610747), were heated to 99°C for 10 min, and were loaded into a pre-cast 12-15% polyacrylamide gel (Bio-Rad, 4561046). Following gel electrophoresis, .22 μm PVDF membranes were used to transfer the proteins via semi dry transfer. Membranes were then blocked in Tris-buffered saline with 0.1% Tween 20 (TBST) containing 2% Bovine serum albumin (BSA) for 1 h. Following blocking, membranes were incubated in polyclonal rabbit anti-LC3C antibody (ProteinTech, 18726-1-AP; 1:1000), anti-UVRAG antibody (Invitrogen, PA5-35213; 1:1000), anti-Beclin 1 rabbit antibody (Cell Signaling Technology, 3495; 1:1000), and anti-ATG14 rabbit antibody (Cell Signaling Technology, 96752; 1:1000), which were detected via anti-rabbit (Cell Signaling Technology, 7076; 1:2000) or anti-mouse HRP-conjugated secondary antibody (Cell Signaling Technology, 7074; 1:2000). The same membranes were stripped and probed with mouse anti-GAPDH (Santa Cruz Biotechnology, sc-47724; 1:1000), as a loading control, and anti-HSp27 mouse antibody (Santa Cruz Biotechnology, sc-13132; 1:1000) for confirmation of HSp27 depletion. These were then detected via anti-mouse HRP-conjugated secondary antibody (Cell Signaling Technology, 7074; 1:2000). Protein bands were viewed via enhanced chemiluminescence (ECL) and band intensities were quantified using NIH ImageJ.

### Far Western Overlay Analysis

5 µg of recombinant LC3C (Origene, TP761768), recombinant LC3B (Sino Biological, 14555-H07E), and recombinant LC3A (Novus Biologicals, NBP1-45308-0.1mg) were resolved by SDS-PAGE as previously in Far Western methodologies [70–72]. The SDS-PAGE resolved protein samples were then transferred to a .22 μm PVDF membrane and were blocked overnight in 5% non-fat milk. The blots were washed and incubated with 10 μg of HSp27 (R&D Systems, 1580HS050) for 1 h. PVDF membranes were probed for HSp27 binding to LC3C, LC3B, or LC3A using a mouse antibody to human HSp27 (Santa Cruz Biotechnology, sc-13132; 1:1000) followed by anti-mouse HRP-conjugated secondary antibody (Cell Signaling Technology, 7074; 1:2000). Protein bands were viewed via enhanced chemiluminescence (ECL) and band intensities were quantified using NIH ImageJ. Control western blots were performed to examine the presence of LC3C, LC3B, or LC3A serve as loading controls using polyclonal rabbit anti-LC3C antibody (ProteinTech, 18726-1-AP; 1:1000), LC3B antibody (Cell Signalling Technology, 3868T; 1:1000), LC3A antibody (Cell Signalling Technology, 12741; 1:1000). Anti-rabbit HRP-conjugated secondary antibody (Cell Signaling Technology, 7076; 1:2000) was then utilized to detect the LC3C, L3CB, or LC3A primary antibody. Antibody cross-reactivity was accounted for via probing non-incubated LC3C blots with mouse anti-HSp27 antibody (Santa Cruz Biotechnology, sc-13132; 1:1000). All Far Western blots were performed at least in duplicate.

### Measurement of Intracellular Glutathione

The levels of glutathione (GSH) of GECs were determined using a luminescence-based GSH/GSSG-Glo Assay kit (Promega, V6611) [41]. This assay is built on the conversion of a luciferin derivative into luciferin, catalyzed by glutathione S-transferase in the presence of GSH. Luminescence signal of each sample was measured with a multi-well plate reader and total levels of GSH was calculated in accordance with the procedure provided. Luminescence for each sample homogenate was converted to GSH concentration based on a standard curve created by serial dilution of a 5 mM GSH standard in the homogenization buffer per the manufacturer’s instructions.

### Co-Immunoprecipitation

Following siRNA transfection, eATP and NAC treatments, and *P. gingivalis* infection, GECs were lysed via extraction buffer, and cell extracts were incubated for 3 h at 4°C with beads pre-coupled overnight with rabbit anti-LC3C (Cell Signaling Technology, 14736, 1:250) antibody. Co-immunoprecipitation was performed with SureBeads™ Protein G Magnetic Beads (BioRad, 1614023) as per the manufacturer’s instructions, and the resulting samples were subjected to 4x Laemmli sample buffer (BioRad, 1610747) and heated to 99°C for 10 min. The eluted proteins were analyzed by western blot as described above.

### Human Biopsy Specimen Specifications

The Medical University of South Carolina’s Institutional Review Board approved the human study protocol and written consent form for tissue collection under the human subjects assurance number, FWA00001888 (with all applicable federal regulations governing the protection of human subjects). Gingival Specimens which were collected from healthy individuals and chronic periodontitis patients (were graded as being in periodontal health, afflicted with moderate periodontitis, or afflicted with severe periodontitis) as we previously described [73] were then anonymized/deidentified and used in this study.

### Immunofluorescence, H&E Staining, and Confocal Microscopy

To initiate staining, the sectioned organotypic rafts were deparaffinized according to the following protocol. In brief, sections were heated at 60°C for 30 min, were incubated in 100% xylene (Fisher Scientific, 016371.K2) for 7 min, and were dipped 20 times in a secondary 100% xylene solution. The samples then underwent rehydration, where they were dipped 20 times in a series of 100%, 95%, and 70% ethanol solutions before they were placed in PBS for 5 min. Slides were then allowed to partially air-dry for 5 min. For H&E staining, samples were stained for 2 min in hematoxylin, differentiated in tap water (15 min) and incubated for 1 min in eosin. For dual antibody staining, rafts were incubated with custom-made 1:200 rabbit anti-*P. gingivalis* ATCC 33277 and mouse anti-HSp27 (Santa Cruz Biotechnology, sc-13132; 1:250) antibodies in PBS for 24 h overnight at 4°C. The samples were then washed briefly in PBS prior to being incubated in 1:500 Alexa Fluor 488 goat anti-rabbit (Invitrogen, A-11008, 1:500) and 1:500 Alexa Fluor 568 donkey anti-mouse (Invitrogen, A10037, 1:500) in PBS for 1h at RT. After secondary antibody incubation, the samples were dipped into PBS 20 times, after which a Vectashield quenching kit (Vector Laboratories, SP-8400-15) was utilized as per the company’s recommendations to limit background fluorescence. After quenching, the samples were stained with 1:500 dilution DAPI for 5 min to visualize cell nuclei, and underwent three 5 min PBS washes. Slides were then air-dried, coverslips were mounted, and imaging was completed the following day using the Zeiss LSM 880 Confocal Microscopy with Airyscan. Control tissue sections without either primary antibodies or secondary antibodies were used in the same manner to further verify the specificity of the antibodies used in this study.

For *in vitro* cell work, primary GECs were plated at a density of 8 × 10^4^ on glass coverslips in four-well plates and cultured until ∼80% confluence. Following siRNA transfection and *P. gingivalis* infection, GECs were fixed with 10% NBF, permeabilized by 0.1% Triton X-100, and stained for 1 h at RT with anti-*P. gingivalis* ATCC 33277 mouse or rabbit antibody (1:1000), anti-HSp27 mouse (Santa Cruz Biotechnology, sc-13132, 1:250) or goat (R&D Systems, AF15801, 1:250) antibody, anti-Beclin 1 rabbit (Cell Signaling Technology, 3495; 1:500 or sheep (R&D Systems, AF5295; 1:250) antibody, ATG14 mouse antibody (Cell Signaling Technology, 96752; 1:250) or anti-LC3C rabbit antibody (ProteinTech, 18726-1-AP: 1:500). After primary antibody incubation, GECs were then washed and incubated in anti-rabbit Alexa Fluor 488 conjugated secondary antibody (Invitrogen, A-11008; 1:1000), anti-mouse Alexa Fluor 488 conjugated secondary antibody (Invitrogen, A-21202; 1:1000), anti-goat Alexa Fluor 405 conjugated secondary antibody (Invitrogen, A48259; 1:250), anti-mouse Alexa Fluor 647 conjugated secondary antibody (Invitrogen, A-31571; 1:1000), anti-mouse Alexa Fluor 405 conjugated secondary antibody (Invitrogen, A48255; 1:250), anti-sheep Alexa Fluor 568 conjugated secondary antibody (Invitrogen, A-21099; 1:1000), anti-mouse Alexa Fluor 568 conjugated secondary antibody (Invitrogen, A10037; 1:1000), or anti-rabbit Alexa Fluor 568 conjugated secondary antibody (Invitrogen, A-11011; 1:1000) at RT for 1 h. GECs were then washed and mounted on glass slides using Vectashield Vibrance mounting media (Vector Laboratories, H-1700-10) Vectashield Vibrance mounting media with DAPI (Vector Laboratories, H-1800-2). Isolated *P. gingivalis* specific autophagosomes were also fixed in 10% NBF for 1 h, and were stained for 1 h at RT with anti-*P. gingivalis* ATCC 33277 rabbit antibody (1:1000), followed by incubation in anti-rabbit Alexa Fluor 488 conjugated secondary antibody (Invitrogen, A-11008; 1:1000). Isolated Vesicles were then incubated in ThiolTracker Violet (Invitrogen, T10095; 1:500) for 30 min, prior to being mounted on glass slides using Vectashield Vibrance Mounting Medium (Vector Laboratories, H-1700-2). Both GECs and Isolated Vesicles were visualized via Zeiss LSM 880 (63x) Confocal Microscopy with Airyscan.

For tissue samples, obtained sections were heated for 30min at 60°C prior to immersion in 100% xylene for 7 min. Following immersion, the samples were then dipped 20 times in a second 100% xylene solution, and underwent a graded series of 20 dips in a 100%, 95%, and 70% ethanol solution respectively. Specimen were then washed for 5 min in PBS, prior to being transferred to a 15mM citrate buffer solution where they were incubated in a pressure cooker on low setting for 15 min to permit for antigen retrieval. After cooking, the slides were allowed to cool at RT for 20 min. The tissues were then exposed to anti-LC3C rabbit primary antibody (Proteintech, 18726-1-AP; 1:500) and either anti-HSP27 mouse primary antibody (Santa Cruz Biotechnology, sc-13132, 1:500) or custom-made mouse anti-*P. gingivalis* primary antibody (1:250) in PBS overnight at 4℃ prior to 20 dips in PBS wash. Secondary antibody staining was completed using Alexa Fluor 568 donkey anti-mouse antibody (Invitrogen, A110037, 1:500), Alexa Fluor 488 goat anti-mouse antibody (Invitrogen, A11001, 1:500), Alexa Fluor 568 donkey anti-rabbit antibody (Invitrogen, A11011, 1:500, and Alexa Fluor 488 goat anti-rabbit antibody (Invitrogen, A11008, 1:500) in PBS for 1h at RT. After secondary antibody incubation, the samples were washed 20 times in PBS, after which the background fluorescence was quenched using a Vectashield quenching kit (Vector Laboratories, SP-8400-15) as per the company’s recommendations. Following quenching, 1:500 DAPI was used to stain nuclear material for 5min at RT, prior to 3 5 min PBS washes. Slides were then air-dried, coverslips were mounted, and imaging was completed the following day. Additionally, control tissue sections without either primary antibodies or secondary antibodies were used in the same manner to further verify the specificity of the antibodies used in this study.

### Transmission Electron Microscopy

Following siRNA transfection and *P. gingivalis* infection, GECs were harvested at 80% confluence, were centrifuged at 1500 x g for 5 minutes, and were placed in 1.5% paraformaldehyde/0.025% glutaraldehyde solution for 1□h, and then in phosphate buffer (PB). GECs were then blocked and permeabilized in 5% Bovine serum albumin (BSA) and 0.1% saponin in PB for 30 min. GECs were then incubated with anti-*P. gingivalis* ATCC 33277 rabbit antibody (1:500) in PB containing 5% BSA and 0.05% saponin for 1 h at RT. Specimen were then washed 3 times for 5 min with PB, prior to incubation with Gold-conjugated secondary antibodies (Transmission Electron Microscopy Sciences, 25100, 1:50) for 1 h. Samples were then washed 3 times for 5 min in PBS, prior to incubation with 2.5% glutaraldehyde. The fixed specimens were dehydrated using a graded ethanol series and were prepared for imaging as described in [74] by the MUSC Transmission Electron Microscopy Core. Ultrathin sections were mounted onto nickel grids. Samples were imaged using a Tecnai BioTwin (FEI Company, Hillsboro, OR, USA) electron microscope operating at 80 kV. Digital images were captured using a 2□K□×□2□K camera (AMT, Danvers, MA, USA).

### GeoData Analysis

mRNA expression data (GEO accession: GSE79705) was obtained from previously collected and examined periodontitis-afflicted and healthy gingival tissues, which is listed on the publicly available GEO repository. The GEO2R program from GEO was then utilized to analyze the microarray expression data and determine the relative levels of HSp27 and LC3C.

### *In silico* molecular docking for the prediction of protein-protein interaction between HSp27 and LC3C

Structures of monomeric full-length wild-type HSP27 (Uniprot: P04792) and LC3C (Uniprot: Q9BXW4) were acquired as models from the AlphaFold database [75]. The algorithm modeled HSP27 with a pLDDT Score (205 residues average) = 68.06 (alpha fold model number: AF-P04792-F1). Accordingly, the pLDDT Score (average of 147 residues) for the LC3C model (alpha fold model number: AF-Q9BXW4-F1) was 79.10. The core secondary structure components of the two proteins were accurately modeled with a pLDDT Score greater than 90. However, the lack of structure in the N- and C-terminal tails increased errors in the models for both proteins. The models were minimized using the Structure Editing module of UCSF Chimera [76]. The default settings used 100 steps of Steepest Descent with a step size of 0.02Å. These were followed by 10 Conjugate Gradient steps using the same step size. All charges and amino acid chemistries were used per the AMBER ff14sb forcefield [77]. PTMs were introduced into the HSP27 using the PyTMs module of PyMOL [78]. Protein-protein docking was done using the Global Range Molecular Matching protocol of GRAMM-X [79,80]. The method uses a Fast Fourier Transform (FFT) based correlation method to find the best surface match between two proteins. Protein-protein complexes are predicted based on a combination of the soft Lennard-Jones potential, the evolutionary conservation of the interface, the statistical choice between residues, the volume of the minimum, the empirical binding free energy, and the atomic contact energy between the proteins. We used a combination of HSP27 and phosphoHSP27 models to obtain their predicted models with LC3C. The GRAMM protocol provided the 10 best-predicted (of the 3000 docked) complexes with the highest empirical binding free energy. All ten complexes (of both regular and phosphorylated HSP27:LC3C) were compared with the Cryo-EM structures of HSP27 (PDBID: 6DV5) to identify the biologically relevant complex configurations. The two complex configurations selected for further analysis had comparable predicted empirical binding free energy of -392 units (HSP27:LC3C) and -395 units (PhosphoHSP27:LC3C). Buried Surface Area/ interaction area between HSP27 and LC3C proteins was assessed by EMBL-PISA [81], and contact maps were generated with MAPIYA [82]. Surface electrostatic potentials of the complexes were mapped using the APBS module of PyMOL [83]. Modeled complexes are available on ModelArchive.org and can be accessed through their IDs: HSP27 LC3C Complex (ma-3foa6) and PhosphoHSP27 LC3C Complex (ma-yy1zt).

### Statistical Analysis

Either one-way ANOVA or two-tailed Student’s t-test were used to evaluate significance. P-values of 0.05 or less were considered to be statistically significant. All experiments were performed at least three separate occasions, in biological triplicate.

**Figure S1.**
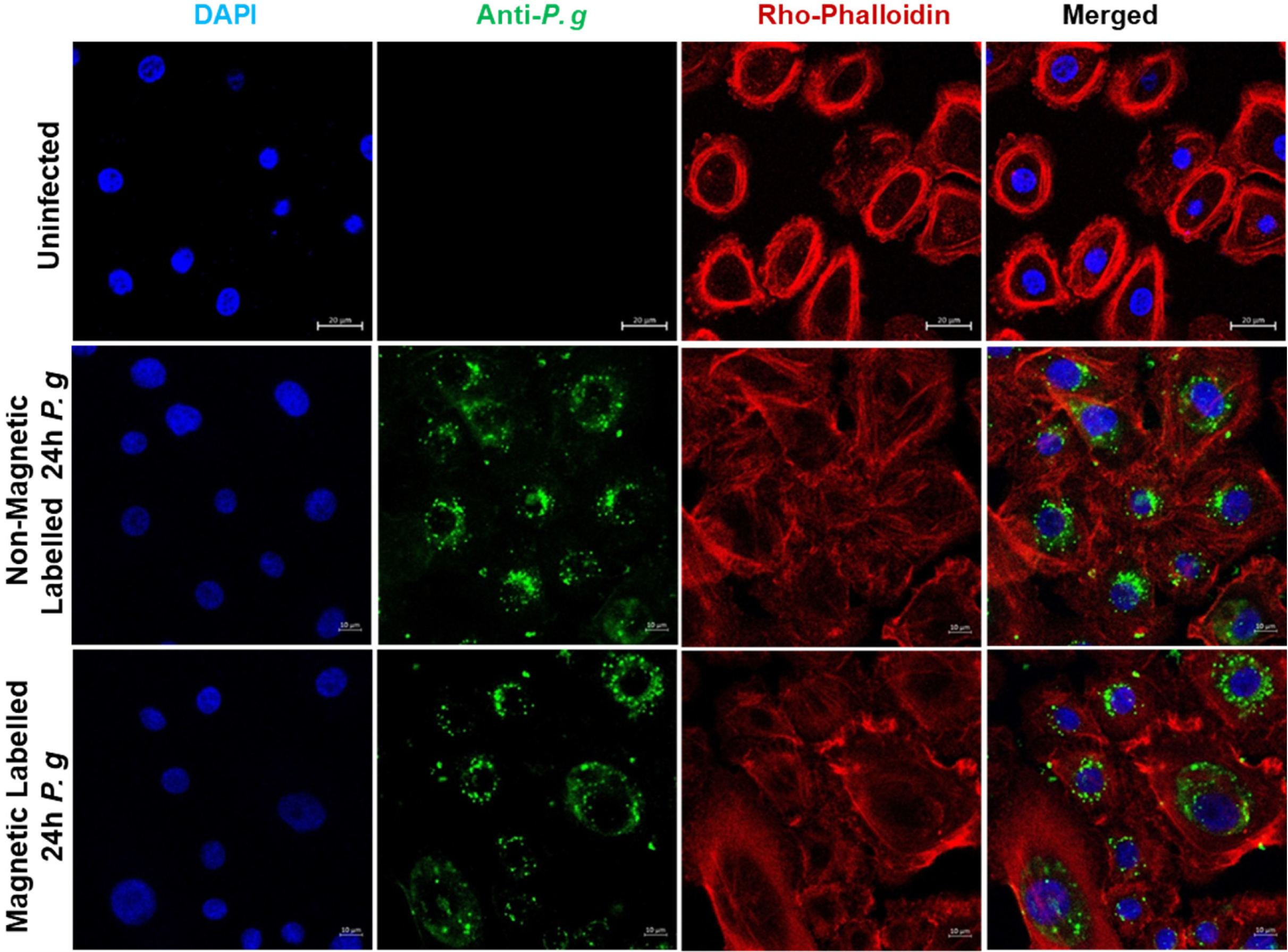
The Magnetic-Labelling Process Does Not Impact *P. gingivalis (P. g)* Infection in GECs. *P. g* was labeled with lipobiotin (5 μM) and were then incubated in MagCellect Streptavidin Ferrofluid. Human Primary GECs were then infected for 24 h. Representative confocal microscopy images of *P. g-*infected GECs at an MOI 100, were taken at 24 h after infection using via Zeiss LSM 880 (63x). *P. g* (rabbit anti-P*. gingivalis*; Alexa 488; green) was detected in the GECs. Actin (red) was stained utilizing Rho-Phallodin.

**Figure S2.**
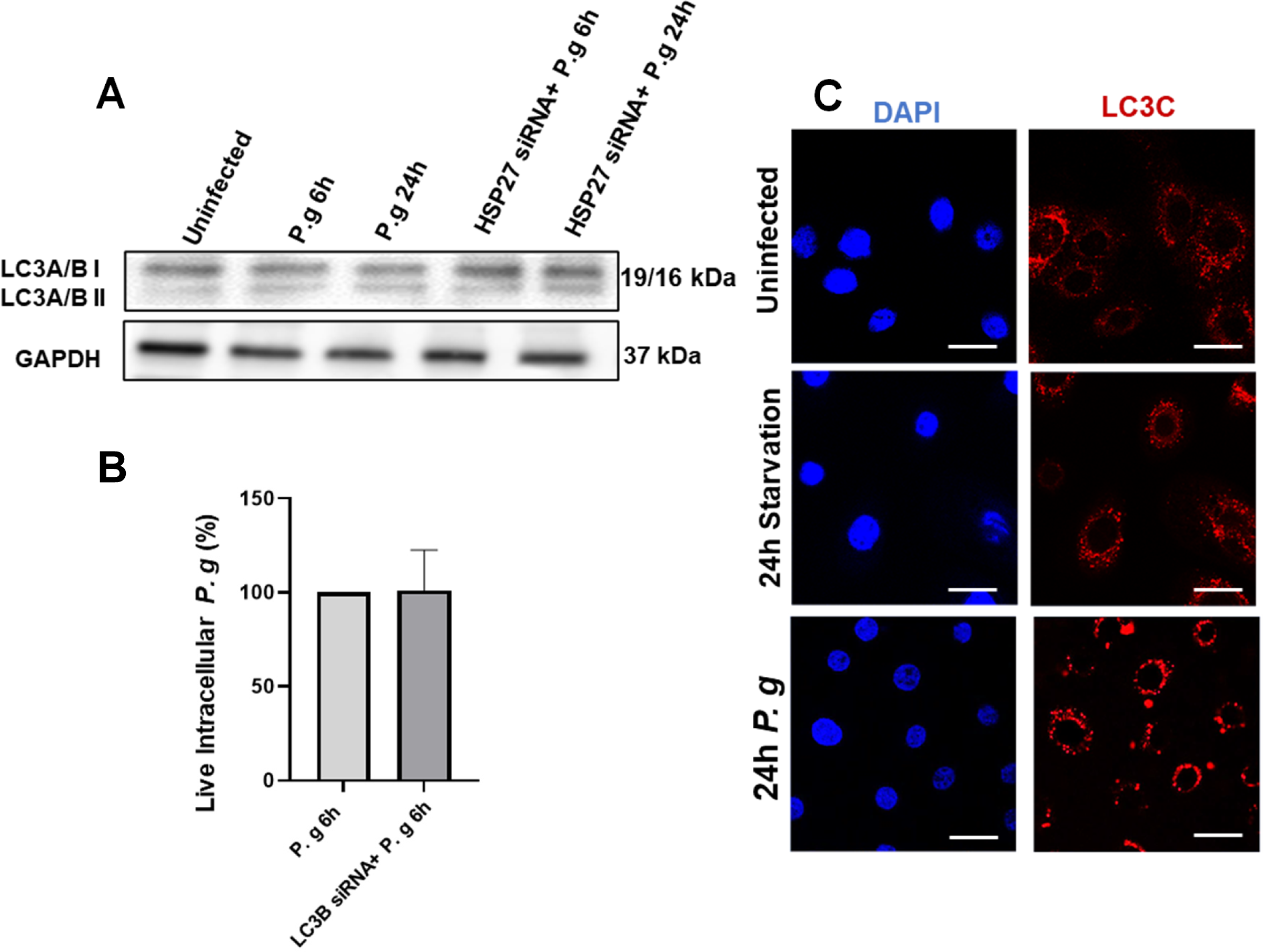
The Autophagic Lifestyle of *P. gingivalis (P. g)* is Highly Characterized by Only the LC3C Isoform of LC3, Which is not Increased During Starvation-Induced Autophagy in GECs. **(A**) The LC3 A/B lipidation results of the same assay provided in Figure 3A**. (B**) GECs were separately treated with LC3B siRNA (100nM) for 48h. *P. g* was added at MOI 100 to GECs for 6 h. Intracellular *P. g* survival after LC3B siRNA depletion was determined using a standard antibiotic protection assay using *P. g-*specific 16S rRNA primers. (**C**) GECs were subjected to starvation conditions in HBSS for 24 h. GECs were then collected and fixed so that immunofluorescence could be performed. GECs were stained for LC3C (rabbit anti-LC3C;Alexa 568; red). GECs were then imaged via confocal microscopy (Super Resolution Zeiss Airyscan LSM 880) at 63x. Western blotting (Not Shown) was utilized to confirm the lack of induction of LC3C I and LC3C II. Data is represented as Mean±SD; n=3; p<0.05 is considered statistically significant (Student two-tailed T-test).

**Figure S3.**
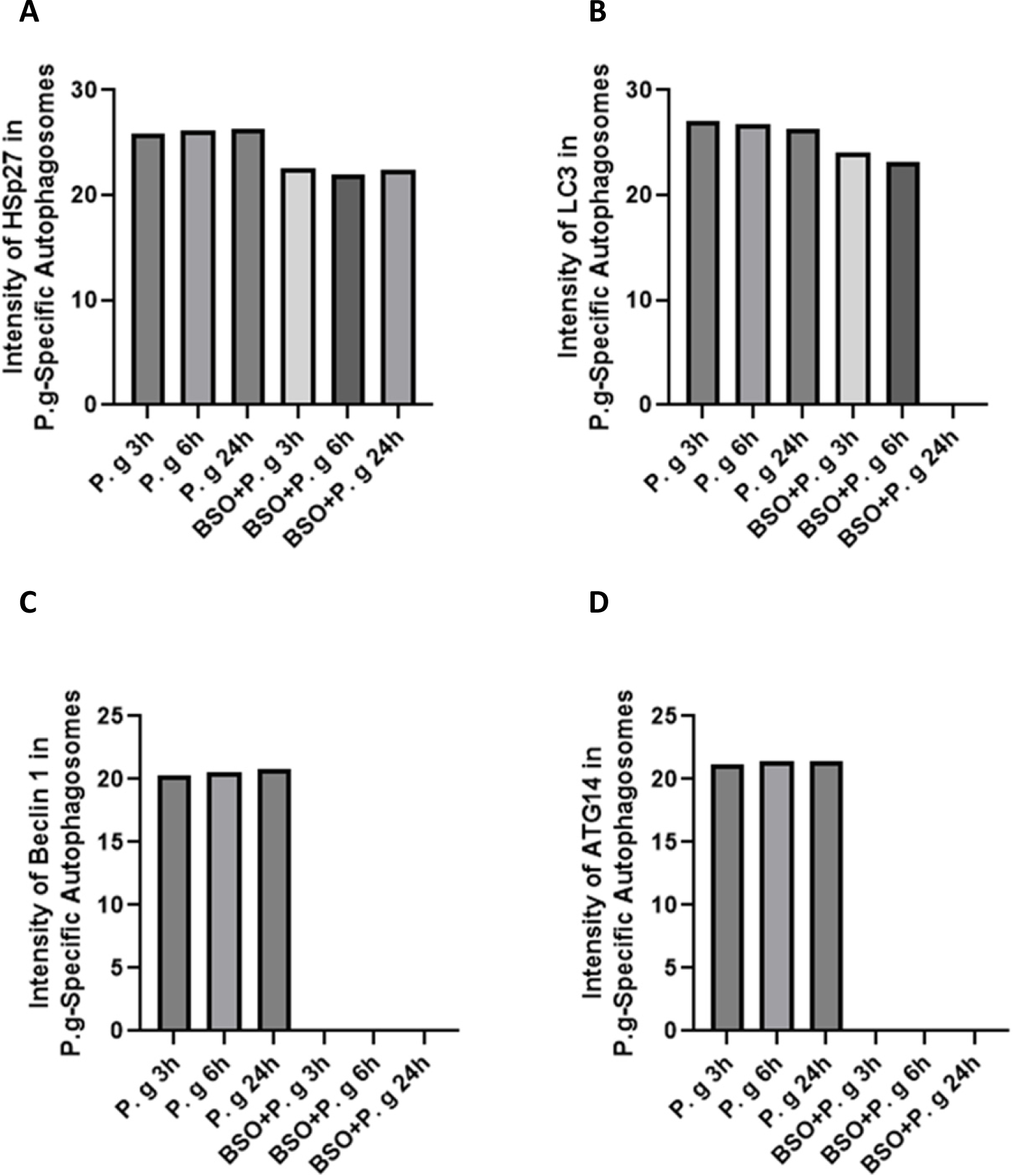
High peak intensity of HSp27, LC3, Beclin 1, and ATG14 in *P. gingivalis*-specific autophagosomes isolated from infected GECs. GECs were pre-treated with buthionine sulfoximine (BSO; 100 nM) for 48 h prior to 3, 6, or 24h of infection with *P. gingivalis*. Following infection, *P. gingivalis-*specific, intact autophagosomes were isolated. The vacuolar proteins were then analyzed by a high resolution and high mass accuracy LC-MS/MS instrument. The label-free quantities (LFQ) of (**A**) HSp27 (Protein ID# P04792), (**B**) LC3 (Protein ID# H3BTL1), (**C**) Beclin 1(Protein ID# O14964), and (**D**) ATG14 (Protein ID#Q6ZNE5)-specific peptides were detected with 95% confidence based on spectrum intensity and Blast search. The assay was performed twice with at least 6-replicates pooled together in each assay.

**Figure S4.**
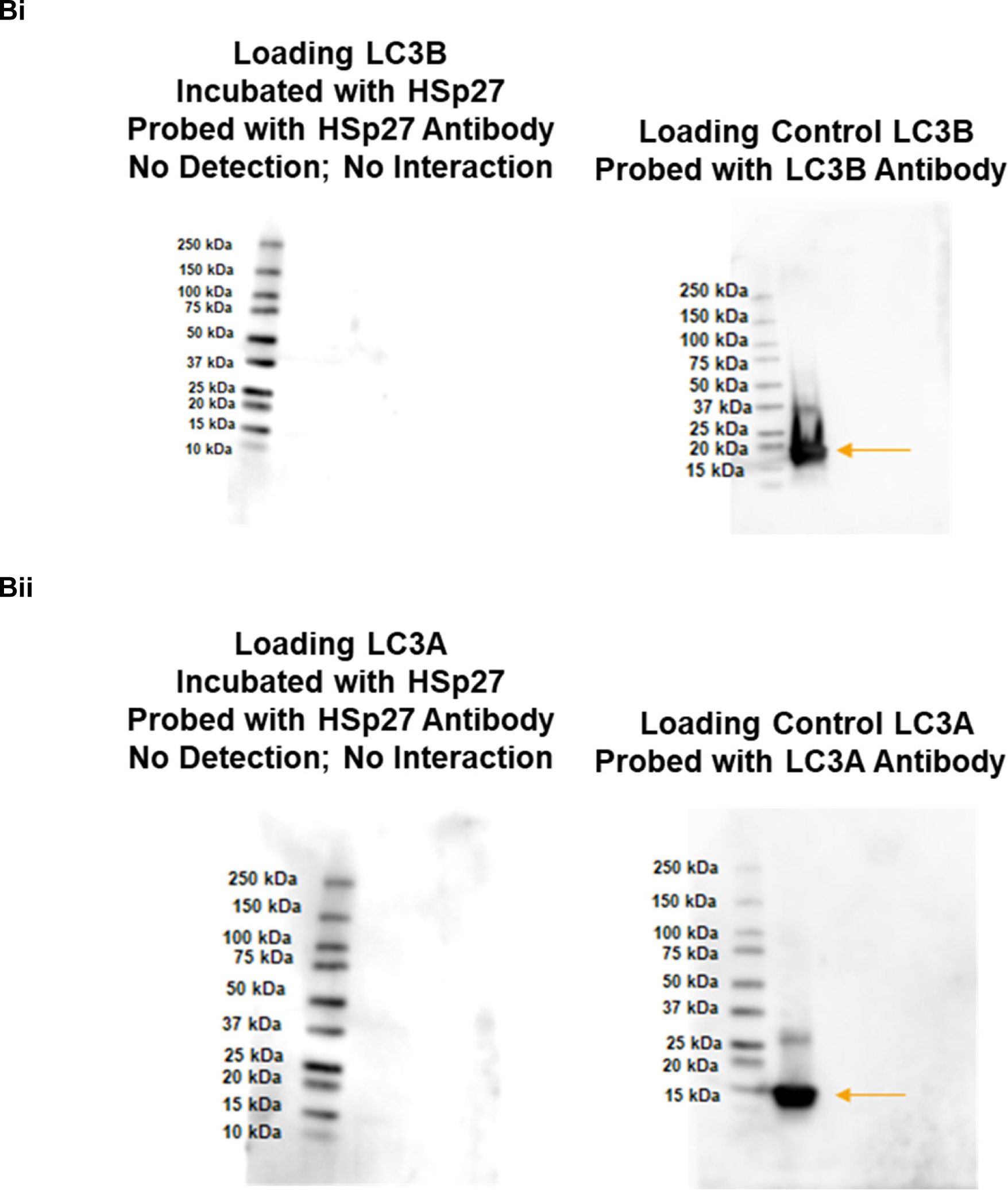
HSp27 does not Interact with either LC3A or LC3B isoforms in the way that it interacts with LC3C. A Far Western approach was implemented. rLC3A or rLC3B were loaded and incubated with 10 ug of rHSp27. Interactions for (**Bi**) LC3A and (**Bii**) LC3B were then detected by probing the rLC3A or rLC3B blot with mouse Anti-Hsp27 antibody.

**Figure S5.**
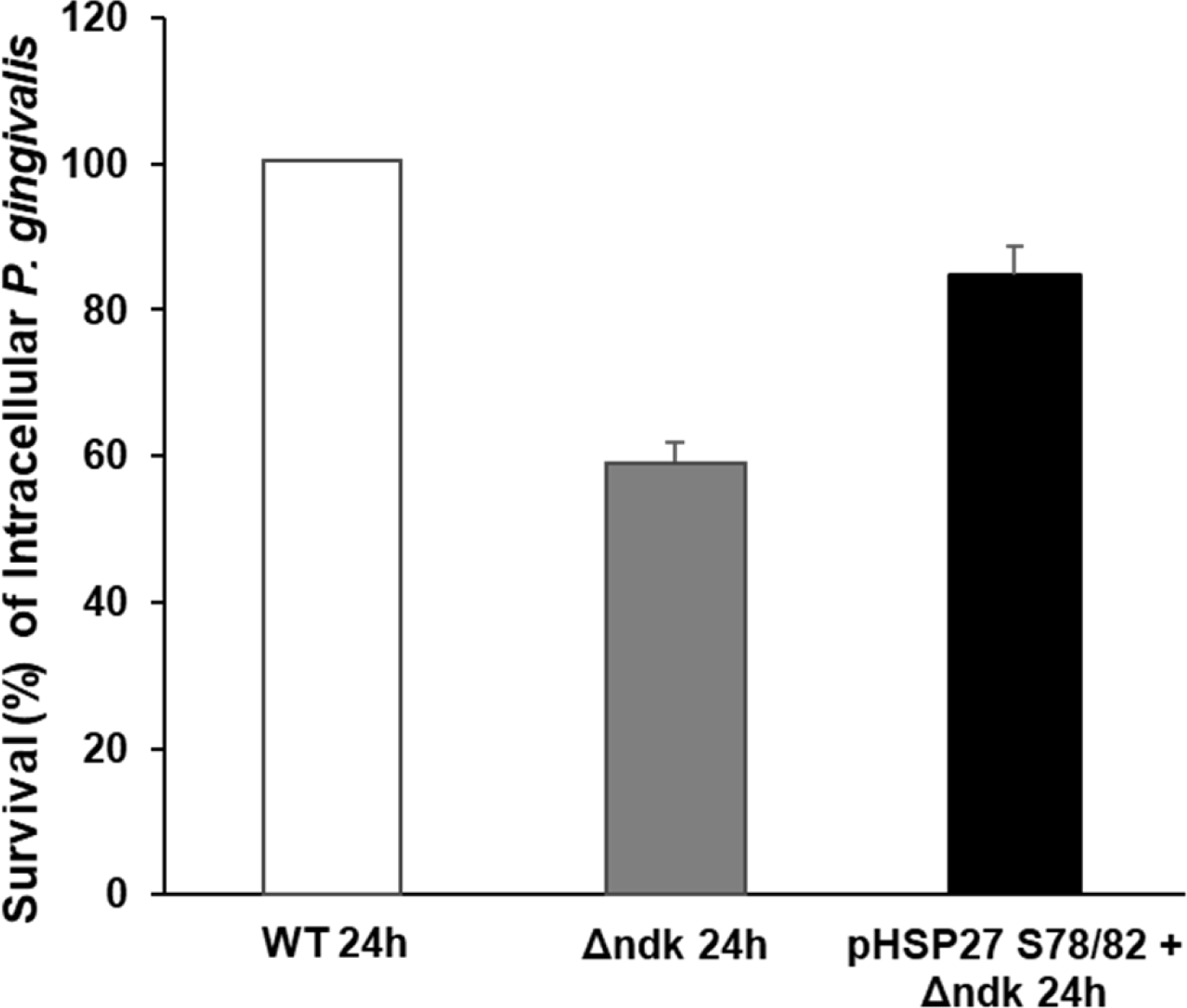
The Intracellular Survival of Δ*ndk P. gingivalis (P. g)* was Restored via Transfection with a constitutively activated HSp27 construct. Briefly, infected GECs were treated with antibiotics and then harvested for qPCR analysis of 16S RNA to quantify intracellular *P. g*. GECs were transfected with pFLAG-CMV2-HSP27-S78D/S82D (1µg) and then infected with WT *P. g* and Δ*Ndk P. g* for 24 hours. Data is represented as Mean±SD; n=3; p<0.05 is considered statistically significant (Student two-tailed T-test).

**Figure S6.**
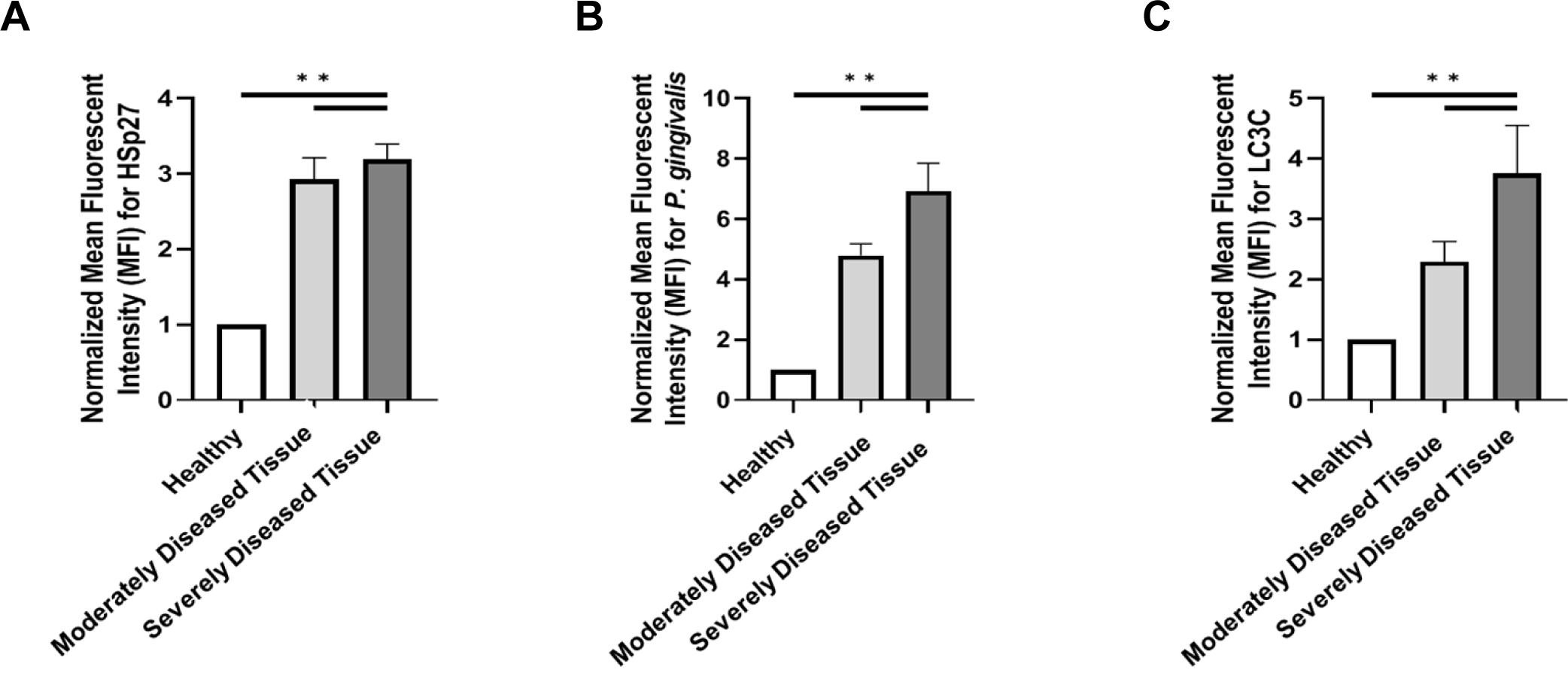
Quantifications of Cross-Sectional Human Ex-Vivo Samples Support High Levels of *P. gingivalis (P. g)*, HSp27, and LC3C in Chronically Diseased Oral Tissues (i.e. Periodontitis). Representative confocal images of gingival biopsy specimens from healthy individuals and periodontitis patients were obtained using the Leica DM6 CS Stellaris 5 Confocal/Multiphoton System so that the mean fluorescence intensity of (**A**) HSp27 (**B**) *P. g* and (**C**) LC3C could be calculated using ImageJ with JACoP Plugin. Data are presented as mean ± SD. Representative images from at least 5 different patients per group were used for quantitative analysis and p<0.05 was considered as statistically significant via Student two-tailed T-test. *p<.05 **p<.005

## AKNOWLEDGEMENTS

This work was primarily supported by funding from the NIH grants R01DE030313, F31DE032273 and T32DE017551. The authors would like to thank Mrs. Sue Newton of the MUSC Transmission Electron Microscopy Core and the MUSC Neuroscience Department for the use of their microscopes. Dr. Shao Yuan of the Hollings Cancer Center Biorepository & Tissue Analysis for his invaluable expertise and help with processing of tissue specimens and interpreting them. MUSC Image facilities were supported in part by Hollings Cancer Center Support Grant (P30 CA138313) and the MUSC Digestive Disease Research Cores Center (P30 DK123704).

